# BRD9 inhibition induces selective radiosensitivity in glioblastoma through MYC pathway modulation

**DOI:** 10.64898/2026.08.10.741731

**Authors:** Nareg Degirmenci, Serdar Aksel Celikkol, Beyza Nur Koseoglu, Adam P Cribbs, Udo Oppermann, Uğur Selek, Tugba Bagci-Onder

## Abstract

Radiotherapy (RT) is a cornerstone of glioblastoma (GBM) treatment, yet therapeutic resistance remains nearly universal due to the rapid activation of stress-adaptive survival programs. Identifying molecular regulators that sustain these adaptive responses may reveal context-dependent vulnerabilities that can be therapeutically exploited. Here, we performed an epigenetic drug screen under low-dose irradiation to identify modifiers of radiotherapy response in glioblastoma. We identify BRD9 inhibition as a priming strategy that selectively enhances irradiation-induced lethality without inducing substantial cytotoxicity under baseline conditions. Mechanistically, BRD9 perturbation delays the resolution of irradiation-induced DNA damage, leading to increased apoptosis following irradiation. This effect is selective for malignant glioblastoma cell lines and patient-derived primary cells, while sparing non-malignant human astrocytes. Transcriptomic profiling reveals that BRD9 inhibition or genetic depletion produces a coordinated, MYC-centered suppression of translational programs, including ribosome biogenesis, rRNA processing, tRNA aminoacylation, and translational initiation. Ectopic MYC expression attenuates BRD9-dependent radiosensitization, functionally linking MYC suppression to the enhanced radiation response. Importantly, analysis of independent glioblastoma patient cohorts reveals a consistent positive association between BRD9 and MYC expression, alongside elevated BRD9 expression in recurrent compared with primary tumors. Together, these findings identify BRD9 as a regulator of MYC-associated translational programs and support its therapeutic targeting as a strategy to enhance radiotherapy efficacy in glioblastoma.

## INTRODUCTION

Glioblastoma (GBM) is the most common and lethal primary brain tumor in adults, characterized by rapid progression and near-universal recurrence despite aggressive multimodal treatment^1^. Current standard of care consists of maximal safe surgical resection followed by concurrent radiotherapy (RT) and temozolomide, with subsequent adjuvant temozolomide cycles^2^. The dismal prognosis reflects the biological complexity of GBM: inter- and intratumoral molecular heterogeneity, efficient DNA damage repair, and the capacity to rapidly reprogram transcriptional states in response to therapeutic stress all conspire to limit durable responses^3,4^. Although RT induces potent DNA double-strand breaks in residual tumor cells, glioblastoma cells activate adaptive stress-response programs that sustain survival and promote recovery from irradiation-induced injury^5^. These adaptive responses, rather than the extent of initial DNA damage, are increasingly recognized as primary drivers of therapeutic resistance and early recurrence^6^. Identifying the molecular regulators that sustain irradiation-induced stress adaptation may therefore reveal context-dependent vulnerabilities that can be exploited to improve radiotherapy efficacy. A growing body of evidence implicates anabolic stress responses, particularly ribosome biogenesis and translational reprogramming, as core components of post-irradiation recovery and stem-cell maintenance in cancer cells^7,8^. In GBM, ribosome biogenesis and protein-synthesis programs have been linked to glioma stem-cell plasticity and to chemo- and radio-resistance, positioning the translational machinery as a candidate therapeutic vulnerability under radiation stress^9,10^.

Epigenetic regulators govern transcriptional plasticity under stress conditions and represent attractive therapeutic targets due to the reversible nature of chromatin modifications^11^. The GBM epigenome is profoundly dysregulated, with aberrant DNA methylation, histone modifications, and chromatin remodeling collectively sustaining malignant transcriptional programs and enabling resistance to both chemotherapy and radiotherapy^12,13^. Irradiation itself induces rapid epigenetic remodeling, including phosphorylation of histone H2AX, alterations in H3K27 methylation, and changes in chromatin accessibility, that modulate DNA damage sensing and repair^14–16^. These observations have motivated extensive efforts to identify epigenetic inhibitors as radiosensitizers, with histone deacetylase (HDAC) inhibitors, Enhancer of Zeste Homolog 2 (EZH2) inhibitors, and Bromodomain and Extraterminal (BET) protein inhibitors emerging as candidate classes^15,17^. However, most approaches have emphasized baseline cytotoxic or anti-proliferative effects rather than context-dependent vulnerabilities that arise specifically under irradiation-induced stress. This distinction matters clinically: an epigenetic inhibitor that selectively enhances radiation sensitivity without substantial single-agent toxicity in normal cells would represent a therapeutically tractable strategy than one that exerts broad cytotoxicity. Identifying epigenetic regulators that are dispensable for basal growth but essential for post-irradiation survival may therefore reveal such exploitable, radiation-specific dependencies.

Bromodomain-containing protein 9 (BRD9) is a defining subunit of the non-canonical BAF (ncBAF) chromatin-remodeling complex, one of three mammalian SWI/SNF subfamilies^18^. BRD9 harbors a bromodomain that preferentially recognizes H3K27ac and related acetylated lysine marks, enabling ncBAF to engage active enhancer and promoter regions^19,20^. Functionally, BRD9 is required for ncBAF complex assembly at target loci, and its inhibition leads to genome-wide dissociation of ncBAF from chromatin^21^. Mutations in SWI/SNF subunits collectively occur in more than 20% of human malignancies, underscoring the oncogenic relevance of chromatin remodeling complexes^22^. While BRD9 itself is not frequently mutated, its expression is elevated or functionally required across multiple cancer contexts. BRD9 has been shown to sustain oncogenic transcriptional programs in acute myeloid leukemia (AML), synovial sarcoma, and multiple myeloma^22–24^. A key mechanistic link has emerged between BRD9 and the MYC oncogene: BRD9 occupies MYC super-enhancer regulatory elements and is required to sustain MYC transcriptional output in cancer cells^23,25^. In multiple myeloma, BRD9 depletion or degradation disrupts ribosome biogenesis gene programs, reduces MYC expression, and impairs protein synthesis machinery^26^, establishing a conserved BRD9–MYC–translational axis across cancer types. Beyond transcription, BRD9 has also been implicated in DNA double-strand break repair: its bromodomain recognizes acetylated RAD54, promoting RAD51–RAD54 complex formation and homologous recombination in non-glial tumor contexts^27^. In GBM specifically, BRD9 has very recently been identified as a tumor-intrinsic determinant of resistance to oncolytic herpes simplex virus type 1 therapy, where its inhibition synergizes with viral antitumor activity through NF-κB–dependent antiviral programs^28^, placing BRD9 within the broader landscape of GBM therapy resistance. Despite these emerging roles, whether BRD9 contributes to GBM cell survival following radiotherapy and whether any radiosensitization is predominantly mediated by direct DNA repair regulation, by transcriptional support of stress-adaptive programs, or by both, has not been systematically investigated.

Here, we identify BRD9 as a regulator of irradiation response in GBM. Through a focused epigenetic drug screen performed under low-dose ionizing radiation, we demonstrate that BRD9 inhibition operates as a priming strategy that selectively enhances irradiation-induced cell death without exerting substantial single-agent cytotoxicity. Transcriptomic profiling across pharmacological and genetic BRD9 perturbation conditions reveals that BRD9 sustains a MYC-dependent transcriptional program. Functional rescue experiments demonstrate that MYC overexpression is sufficient to attenuate BRD9-dependent radiosensitization and restore translational gene outputs, establishing MYC as a causal mediator of the BRD9–radiation axis. Finally, the BRD9–MYC translational program is conserved across independent patient cohorts, supporting the clinical relevance of this axis. Together, these findings define BRD9 as a stress-adaptive, non-oncogene dependency in GBM and identify its inhibition as a strategy to disrupt MYC-dependent translational programs and enhance radiotherapy response.

## MATERIALS & METHODS

### Cell Culture

Human glioblastoma cell lines U373 and U87-MG and human embryonic kidney cells (HEK293T; used for lentiviral particle production) were obtained from the American Type Culture Collection (ATCC, USA). Immortalized normal human astrocytes (I-NHA) were kindly provided by Tim Chan (Memorial Sloan Kettering Cancer Center, MSKCC). All cell lines were cultured in Dulbecco’s Modified Eagle Medium (DMEM; Gibco, USA) supplemented with 10% fetal bovine serum (FBS; Gibco, USA) and 1% penicillin–streptomycin (Gibco, USA). Cells were maintained at 37 °C in a humidified incubator with 5% CO₂. All cell lines were routinely tested for mycoplasma contamination and confirmed to be mycoplasma-free throughout the course of the study. GBM8 (MGG8) cell line was established by Dr. Hiroaki Wakimoto from patient-derived models, at Massachusetts General Hospital (MGH)^29^. GBM8 cells were cultured as neurospheres in GBM/EF medium consisting of Neurobasal medium (Gibco, USA) with 7.5 ml L-Glutamine, 1X B-27 supplement (Gibco, USA), 0.5X N-2 supplement (Gibco, USA), 0.5 ml heparin solution (0.2%, Stemcell Technologies, Canada), 0.5% Pen/Strep, FGF (20 ng/ml) and EGF (20 ng/ml).

### Chemicals and reagents

The epigenetic drug library comprising 124 compounds was kindly provided by Dr. Udo Oppermann (University of Oxford, UK). The full composition of the library is listed in **Supplementary Table 1**. I-BRD9 was purchased from Selleckchem (Cat. #S7835). The BRD7/BRD9 PROTAC degrader VZ185, BRD7/BRD9 inhibitor BI-7273, and BRD9-selective inhibitor BI-9564 were obtained through the opnMe platform (Boehringer Ingelheim).

### Irradiation experiments

All irradiation experiments were performed using an XStrahl CIX2 Cabinet X-Ray Irradiator located at Koç University Center of Translational Medicine Animal Research Facility (KUARF). Cell-seeded plates were placed in the irradiator cabin and exposed to radiation at 195 kV - 10 mA to deliver the indicated doses.

### Epigenetic drug screen

An epigenetic probe library consisting of well-characterized inhibitors targeting chromatin-modifying and chromatin-reading proteins was assembled from the Structural Genomics Consortium (SGC) and other sources. This collection and related chemical-probe libraries have been successfully employed in epigenetic vulnerability screens across diverse cancer models^30,31^. U373 cells were seeded at 1,000 cells per well in round-bottom 96-well plates. The following day, cells were treated with epigenetic compounds at one-tenth of their working concentration (0.1X). On day 3, cells were exposed to a single dose of ionizing radiation (IR-4 Gy). After 24 hours, cells were trypsinized and 10% of the collected cells were reseeded into black 96-well plates for viability assessment using the CellTiter-Glo (CTG) assay. Cell viability was measured on day 7 according to the manufacturer’s instructions. The mean viability of DMSO-treated irradiated controls was calculated (68.59% ± 5.06%, mean ± SEM). Radiosensitization was quantified as the difference between drug-only and drug + IR conditions. The screen was performed with triplicates, with mean viability values used for hit ranking. Hits were defined operationally as compounds whose mean viability under combination treatment fell at or below the mean viability of DMSO + IR controls, and library compounds annotated as inactive controls were excluded from the hit list.

### Clonogenic survival assay

Cells were seeded at 200–250 cells per well in 12-well plates in triplicate and treated with the indicated inhibitors for 72 hours. Cells were then irradiated with single doses of 2, 4, 6, or 8 Gy and incubated for 10–14 days. For schedule comparisons, I-BRD9 was added either 72 h before IR (pre-treatment), at the same time as IR (co-treatment), or 24 h after IR (post-treatment); cells in all three arms were processed on the same timeline. For CRISPR/Cas9-edited cells, clonogenic assays were performed six days post-transduction, followed by irradiation the next day. Colonies were fixed, stained, and quantified as previously described ^32^. Colony area density was quantified using Adobe Photoshop CC 2019.

### MTT assay

Cells were seeded at 750 cells per well in 96-well plates, treated with inhibitors on day 1, and irradiated on day 3. On day 5, MTT reagent (3 mg/mL in PBS) was added and incubated for 4 hours at 37 °C. After removal of the medium, formazan crystals were dissolved in DMSO and absorbance was measured at 570 nm using a Synergy H1 microplate reader (BioTek).

### Cell-Titer Glo (CTG) assay

Cells were seeded as 1000 cells/well to 96-well black culture plates and next day treated with the selected inhibitors. After 3 days, all plates were irradiated. On day 7, cell viability was assessed according to the manufacturer’s instruction using a plate reader (BioTek’s Synergy H1, Winooski, VT, USA).

### Spheroid assay

Primary patient-derived cells were seeded 1,000 cells per well to generate spheroids, allowed to self-assemble, and monitored for spheroid formation over 5 days. Following spheroid establishment, cells were treated with I-BRD9 at 1, 2, 5, and 10 µM. After a 3-day drug incubation period, spheroids were exposed to IR at 2 Gy and 4 Gy. Spheroid growth and morphology were monitored for several days post-treatment; on day 10, spheroid size was quantified from brightfield images by image-based measurement, and cell viability was assessed using the CellTiter-Glo (CTG) assay according to the manufacturer’s instructions.

### Immunofluorescence staining

Cells were seeded on coverslips in 24-well plates at 10,000 cells/well. Following treatment, cells were fixed in 4% paraformaldehyde for 5 minutes, permeabilized with 0.1% Triton X-100, and blocked with SuperBlock IHC Blocking Solution. Primary antibodies were incubated overnight at 4 °C, followed by secondary antibodies for 1 hour at room temperature in the dark. Coverslips were mounted with DAPI-containing mounting medium. Images were acquired using a Leica DMI8 inverted microscope equipped at 40× magnification. At least 120 nuclei were quantified per condition across three independent replicates, and a mean nuclear signal intensity was calculated with Fiji. Antibodies are listed in **Supplementary Table 2**.

### Western blotting

Protein extraction and immunoblotting were performed as previously described^33^. Antibodies are listed in **Supplementary Table 2.**

### Quantitative Real-Time PCR (qRT-PCR)

RNA isolation and cDNA synthesis were performed as previously described^5^. Primer sequences are listed in **Supplementary Table 3.**

### RNA-sequencing and analysis

Total RNA was isolated using the NucleoSpin RNA Isolation Kit (Macherey-Nagel). Libraries from U87-MG BRD9 knockout samples were sequenced at Oxford University NDORMS using a NextSeq 500 instrument, generating ∼24 million single-end reads per sample. Libraries from U373 and U87-MG I-BRD9-treated samples and U373 BRD9 knockout samples were sequenced using the BGISEQ-500 platform. Reads were processed on the Genialis Expressions platform using the alignment-free Salmon workflow with default parameters unless stated. Adapters and low-quality bases were trimmed with BBDuk. Transcripts were quantified with Salmon v1.2.1 in mapping-based mode against a decoy-aware index built from

Ensembl release 100 human cDNA (primary assembly), extended with ERCC/SIRV spike-in sequences and the full genome as decoy, using --libType A, --validateMappings, --seqBias, --gcBias, and --rangeFactorizationBins 4. Transcript-level estimates were summarized to gene-level counts and TPM with tximport. Differential expression was assessed with DESeq2 (Wald test), with genes at adjusted p < 0.05 and |log₂ fold-change| > 1 considered significant; pathway enrichment used fgsea against MSigDB Hallmark and curated gene sets. Three biological replicates were sequenced per condition.

### CRISPR mediated knock-out

sgRNAs were designed to target exonic regions of human BRD9 using the ChopChop gRNA design tool^34^. Oligonucleotides were annealed and cloned into pLentiGuide-Puro (Addgene #117986) as described^35^. sgRNA sequences were listed in **Supplementary Table 4.**

### Dual H2AX activation assay

Cells were harvested from 6-well plates following the corresponding treatment, and H2AX activation was quantified with The Muse® H2AX Activation Dual Detection Kit (MCH200101) according to the manufacturer’s protocol.

### Annexin/PI staining

Annexin/PI staining was performed with BD Pharmingen™ FITC Annexin V Apoptosis Detection Kit I (#556547) according to the manufacturer’s protocol. The staining was performed as previously described^36^. Three independent replicates were analyzed for each condition.

### Publicly available data analysis

Vital status data were downloaded via cBioPortal^37^ using the Glioblastoma (CPTAC, Cell 2021), Brain Tumor PDXs (Mayo Clinic, Clin Cancer Res 2020), Glioblastoma (TCGA, Cell 2013) and Diffuse Glioma (GLASS Consortium, Nature 2019) datasets, and also Gliovis TCGA GBM microarray dataset (HG-UG133A platform)^38^. Primary versus recurrent BRD9 and MYC expression was compared in the TCGA GBM Affymetrix HG-U133A microarray cohort using a two-tailed Mann–Whitney test. For BRD9–MYC co-expression analyses across the GLASS, CPTAC, TCGA GBM, and Mayo PDX cohorts, mRNA expression matrices were retrieved through cBioPortal and Spearman rank correlation coefficients between BRD9 and MYC were computed with associated p-values; the same approach was applied genome-wide to derive the per-gene Spearman correlations with BRD9 and with MYC used to construct the bubble and scatter plots. BRD9 gene-effect (Chronos) scores across cancer lineages were obtained from the Cancer Dependency Map (DepMap) portal (depmap.org; release 26Q1) and visualized as lineage-resolved distributions, with rhabdoid and synovial sarcoma lines shown as a positive-dependency reference and CNS/glioma lineages highlighted.

### ChIP Atlas analysis

For chromatin occupancy analysis at the promoters of the 31 conserved BRD9-dependent genes, the ChIP-Atlas Enrichment Analysis tool was queried using gene symbols as input against all publicly deposited ChIP-seq datasets in the neural cell type class (H. sapiens, hg38 assembly, MACS2 score threshold ≥ 50). Enrichment was assessed by Fisher’s exact test and expressed as fold enrichment over background.

### Statistical analysis

All normalizations were performed on non-irradiated or untreated samples denoted as 100% using GraphPad Prism version 9.0 (USA) and Microsoft Excel 2018. Data are presented as mean ± SEM unless otherwise stated. Significance analysis was performed with student’s t-test and two-way ANOVA (n.s denote not significant, for p-values, *, ** and *** denote p < 0.05, p < 0.01 and p < 0.001 respectively, two-tailed Student’s t-test).

## RESULTS

### BRD9 inhibitors are identified as radiosensitizers through an epigenetic screen in glioblastoma

To systematically identify epigenetic regulators that modulate the radiotherapy response of GBM, we performed a focused chemical screen in U373 cells using a library of 124 epigenetic probes spanning bromodomain, histone methyltransferase, HDAC, histone demethylase, and related target classes **(Figure 1A).** Compounds were applied at one-tenth of their working concentration on day 1 (0.1x), cells were irradiated with a single 4 Gy dose on day 4, and viability was assessed by CellTiter-Glo on day 8 **(Figure 1B).** By ranking compounds according to their ability to reduce viability specifically in combination with irradiation rather than as single agents, the screen was designed to enrich for context-dependent radiosensitizers over baseline cytotoxic agents **(Figure 1C).** Heatmap visualization of the top 20 radiosensitizing compounds confirmed that most hits displayed minimal single-agent cytotoxicity yet markedly reduced viability under drug-plus-irradiation conditions **(Supplementary Figure S1A).** Bromodomain inhibitors emerged as the most prominent class among the screen hits **(Figure 1D),** with BRD9-targeting compounds, LP99, TP-472, and I-BRD9 consistently ranking among the strongest radiosensitizers.

**Figure 1.**
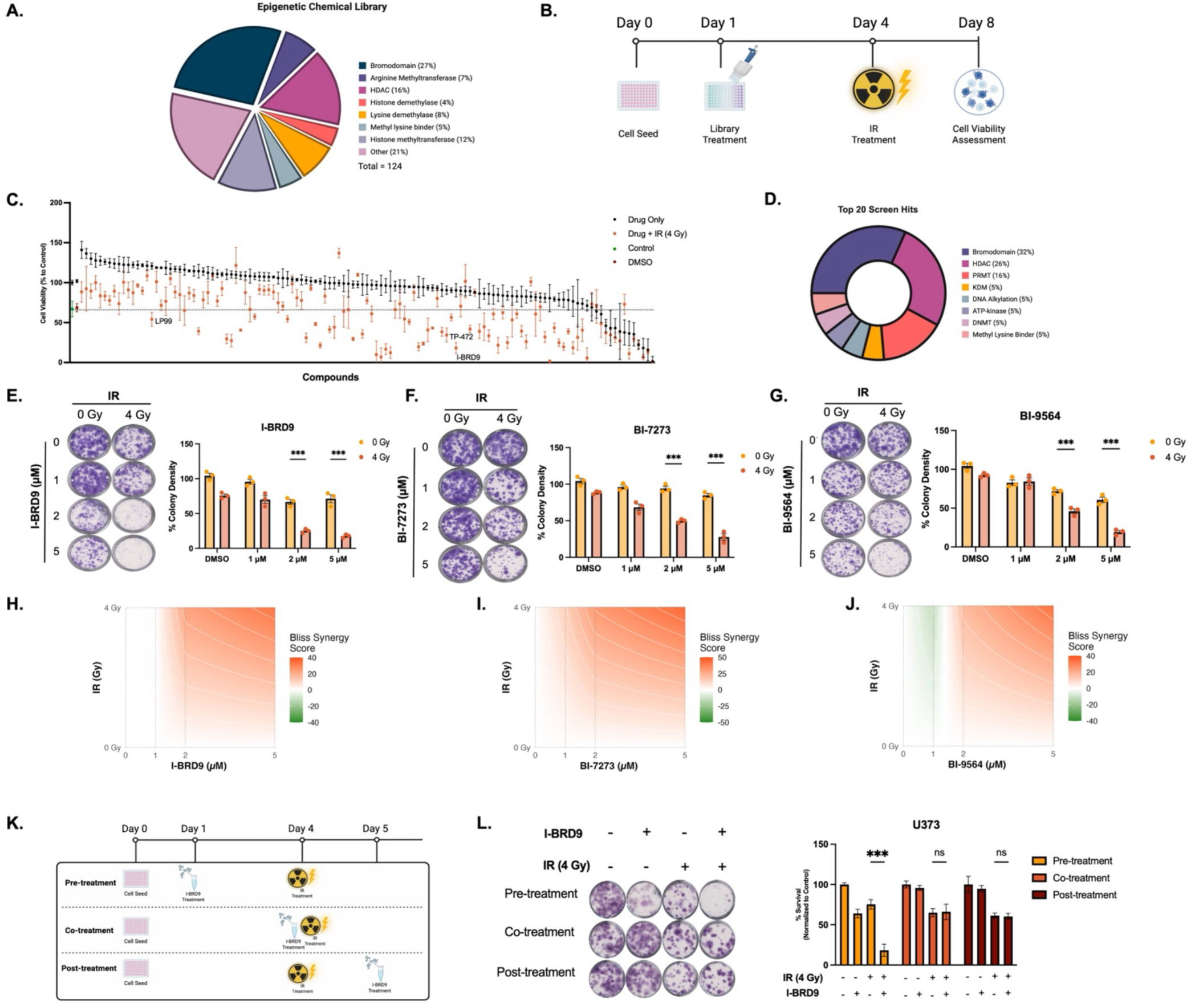
Epigenetic drug screening identifies BRD9 inhibition as a priming strategy for radiation response in glioblastoma. **A.** Composition of the 124-compound epigenetic probe library used for the screen, grouped by target class. Generated in BioRender. **B.** Schematic of the screening workflow. U373 glioblastoma cells were seeded on day 0, treated with individual library compounds at one-tenth of their working concentration on day 1, irradiated with a single 4 Gy dose on day 4, and assessed for viability on day 8. Generated in BioRender. **C.** Ranked cell viability (% of untreated control, mean ± SEM) of the 124 epigenetic compounds in U373 cells under drug-only (black) or drug + ionizing radiation (4 Gy; magenta) conditions. Compounds are ordered by their viability in the combination arm. The dashed line indicates the mean viability of DMSO + IR controls, used as the hit-calling threshold. **D.** Distribution of classes of epigenetic modifier hits from the screen classes. Generated in BioRender. **E,F,G.** Representative clonogenic survival images (left) and colony-density quantification (right) for U373 cells treated with increasing concentrations of three structurally distinct BRD9 inhibitors (E) I-BRD9, (F) BI-7273, and (G) BI-9564 in the absence or presence of IR. Colony density was normalized to DMSO-treated, non-irradiated controls. **H, I, J.** Two-dimensional Bliss synergy landscapes for the drug × radiation combinations shown in panels E, F, and G, computed with SynergyFinder from the corresponding viability matrices. Red indicates synergy (Bliss score > 0) and green indicates antagonism (Bliss score < 0). **K.** Schematic representation of BRD9 inhibitor treatment schedules relative to IR, illustrating pre-treatment, co-treatment, and post-treatment strategies used in clonogenic survival assays. Generated in BioRender**. L.** Representative clonogenic survival images and quantification of normalized colony density for U373 cells treated according to each schedule in panel K with I-BRD9 (2 µM) and/or 4 Gy irradiation, as indicated below the axis.

To confirm this on-target BRD9 dependency rather than assay-specific or off-target activity, we first re-tested the BRD9-targeting screen hits in clonogenic survival assays: LP99, TP-472, and I-BRD9, together with the selective BRD9 bromodomain inhibitor BI-9564. Each radiosensitized U373 cells, causing a marked loss of clonogenic survival in combination with irradiation while having little effect as single agents (**Supplementary Figure S1B**). Since these probes differ in potency and chemotype, we then focused on the well-characterized, selective inhibitor I-BRD9 for detailed mechanistic studies and independently validated BRD9 dependency with two additional structurally distinct, selective BRD9 bromodomain inhibitors, BI-7273 and BI-9564. In clonogenic survival assays, all three inhibitors reduced colony-forming capacity in a concentration-dependent manner when combined with 4 Gy irradiation, while exerting only modest effects in the absence of irradiation (**Figure 1E–G**). To quantify the drug–radiation interaction, we computed two-dimensional Bliss synergy landscapes across the inhibitor-concentration and irradiation-dose ranges; all three inhibitors produced positive Bliss synergy scores, indicating that BRD9 inhibition and irradiation act synergistically rather than additively (**Figure 1H–J**). BRD9 dependency was further supported by the BRD7/BRD9 PROTAC degrader VZ185, which depleted BRD9 protein within 6 hours and enhanced radiosensitization across increasing radiation doses (**Supplementary Figure S2A-C**).

To determine whether the timing of drug application and irradiation influences overall efficacy, we compared three treatment schedules, pre-treatment, co-treatment, and post-treatment relative to irradiation **(Figure 1K)**. Pre-treatment with I-BRD9 before irradiation produced the most pronounced reduction in clonogenic survival, significantly outperforming co- and post-treatment, which were not significantly different from irradiation alone in U373 and U87-MG cells **(Figure 1L, Supplementary Figure S2D).** This priming behaviour is consistent with BRD9 inhibition establishing a vulnerable state before irradiation that the post-irradiation window does not recapitulate. Together, these findings establish BRD9 inhibition as a reproducible, synergistic priming strategy that enhances irradiation-induced lethality in GBM cells.

### BRD9 inhibition enhances irradiation-induced apoptosis and delays resolution of irradiation-induced DNA damage

To define the cellular basis of I-BRD9-dependent radiosensitization, we examined apoptosis and DNA damage response after combined BRD9 inhibition and irradiation in U373 cells. Annexin V/PI flow cytometry revealed a marked increase in both early- and late-apoptotic populations in cells receiving combination treatment relative to either treatment alone (**Figure 2A,B**), indicating that BRD9 inhibition potentiates irradiation-induced apoptotic cell death. To ask whether altered DNA-damage kinetics accompany this effect, we monitored γH2AX and 53BP1 at 30 minutes, 2 hours, and 24 hours after irradiation (**Figure 2C**). Immunofluorescence imaging of γH2AX and 53BP1 foci (**Figure 2D**) showed that combination-treated cells retained a high proportion of foci-positive cells at 24 hours, whereas foci largely resolved in irradiation-only cells (**Figure 2E,F**). Consistently, flow-cytometric quantification of γH2AX showed that irradiation alone induced a transient response that declined by 24 hours, whereas combination-treated cells sustained elevated γH2AX at 24 hours despite a lower initial signal, indicative of delayed damage resolution rather than a greater initial damage burden (**Figure 2G,H**). The same enhancement of apoptosis was observed in U87-MG cells (**Supplementary Figure S3**), demonstrating that increased apoptosis and delayed DNA-damage resolution are conserved consequences of BRD9 perturbation under irradiation stress across GBM models.

**Figure 2.**
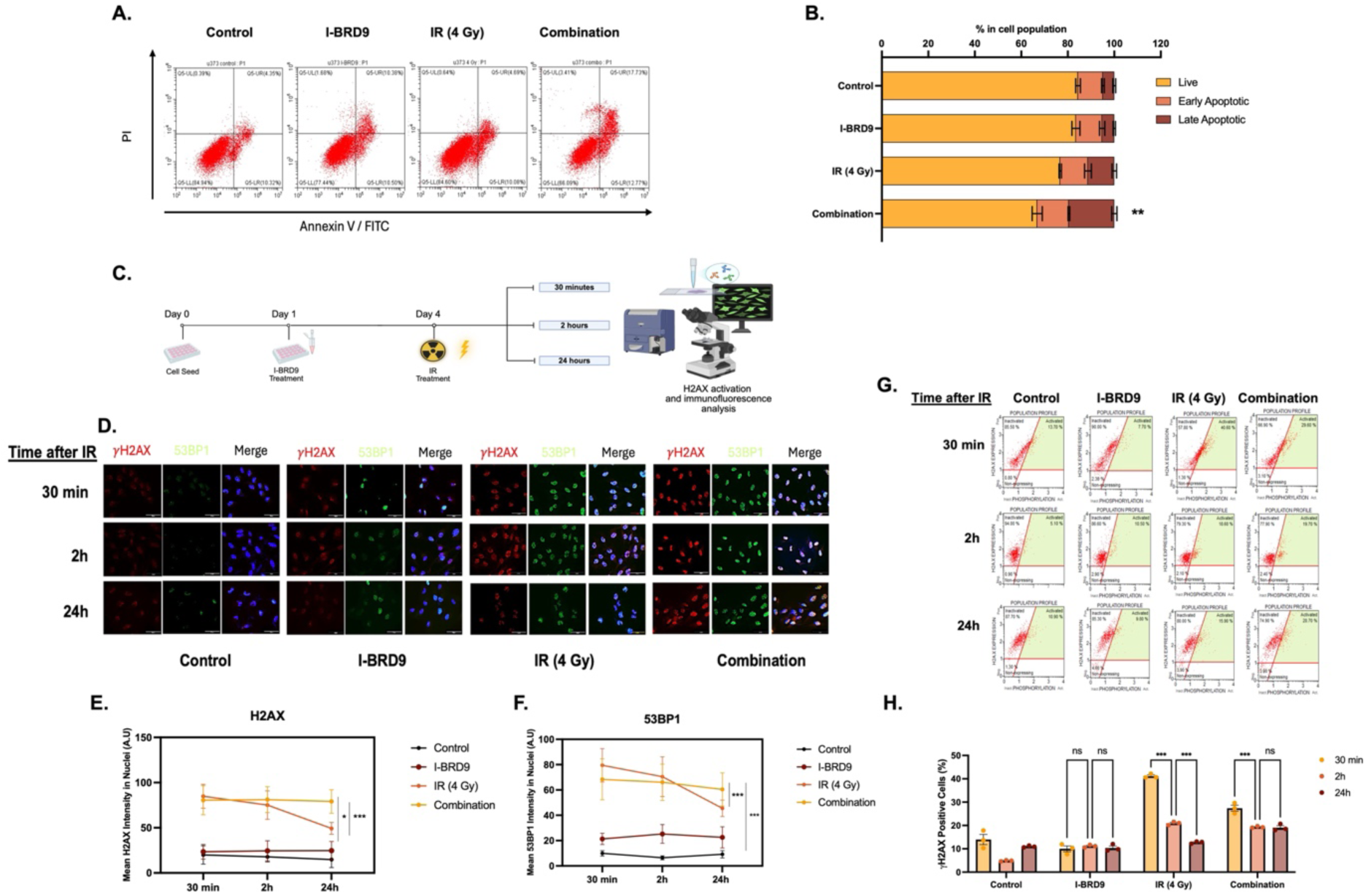
BRD9 inhibition enhances radiation-induced apoptosis and delays DNA damage resolution in glioblastoma cells. **A.** Representative Annexin V/PI flow cytometry plots of U373 cells 48 hours after treatment with DMSO, I-BRD9 (2 µM), IR (4 Gy), or the combination of I-BRD9 and IR. **B.** Quantification of live, early apoptotic, and late apoptotic cell populations following the indicated treatments. **C.** Schematic of the immunofluorescence and H2AX activation assay time-course experiment. U373 cells were seeded on day 0, treated with I-BRD9 on day 1, irradiated on day 4, and fixed at 30 min, 2 h, and 24 h post-irradiation for confocal imaging. Generated in BioRender. **D.** Representative immunofluorescence images of γH2AX (red) and 53BP1 (green) foci in U373 cells at 30 minutes, 2 h, and 24 h following treatment. Nuclei were counterstained with DAPI (blue). Scale bar, 20 µm. **E,F**. Quantification of mean nuclear (E) γH2AX and (F) 53BP1 immunofluorescence intensity over time. Each point represents the mean ± SD of all analysed nuclei from one representative experiment (120 nuclei per condition/timepoint), measured objectively by automated nuclear segmentation (DAPI) and per-nucleus intensity quantification under identical acquisition settings. **G.** Representative Muse H2A.X Activation Dual Detection flow-cytometry profiles of U373 cells at 30 min, 2 h, and 24 h after irradiation for each treatment arm. The activated (γH2AX-positive) gate is shown in green with the corresponding percentage. **H.** Quantification of γH2AX-positive cells across the time course is shown below; data are mean ± SEM of three independent biological replicates

### BRD9-dependent radiosensitization is selective for malignant glioblastoma cells

To test whether BRD9-mediated radiosensitization is selective for malignant cells, we compared BRD9 perturbation in two GBM lines and immortalized normal human astrocytes (I-NHA). Dose– response analysis showed that I-BRD9 had minimal impact on I-NHA viability at any concentration, with or without irradiation, whereas both GBM lines exhibited significant, dose-dependent loss of cell viability upon combined I-BRD9 and irradiation (**Figure 3A**). In clonogenic assays, combination treatment significantly reduced colony formation in U373 and U87-MG cells but not in I-NHA (**Figure 3B**), defining a selective therapeutic window. Genetic validation using two independent BRD9-targeting sgRNAs (BRD9-g1, BRD9-g2), confirmed at the protein level (**Figure 3C**), recapitulated this pattern: BRD9 loss caused a modest reduction in baseline proliferation (**Supplementary Figure S4**), but markedly reduced post-irradiation clonogenic survival in both GBM lines while leaving I-NHA largely unaffected (**Figure 3E,F**).

**Figure 3.**
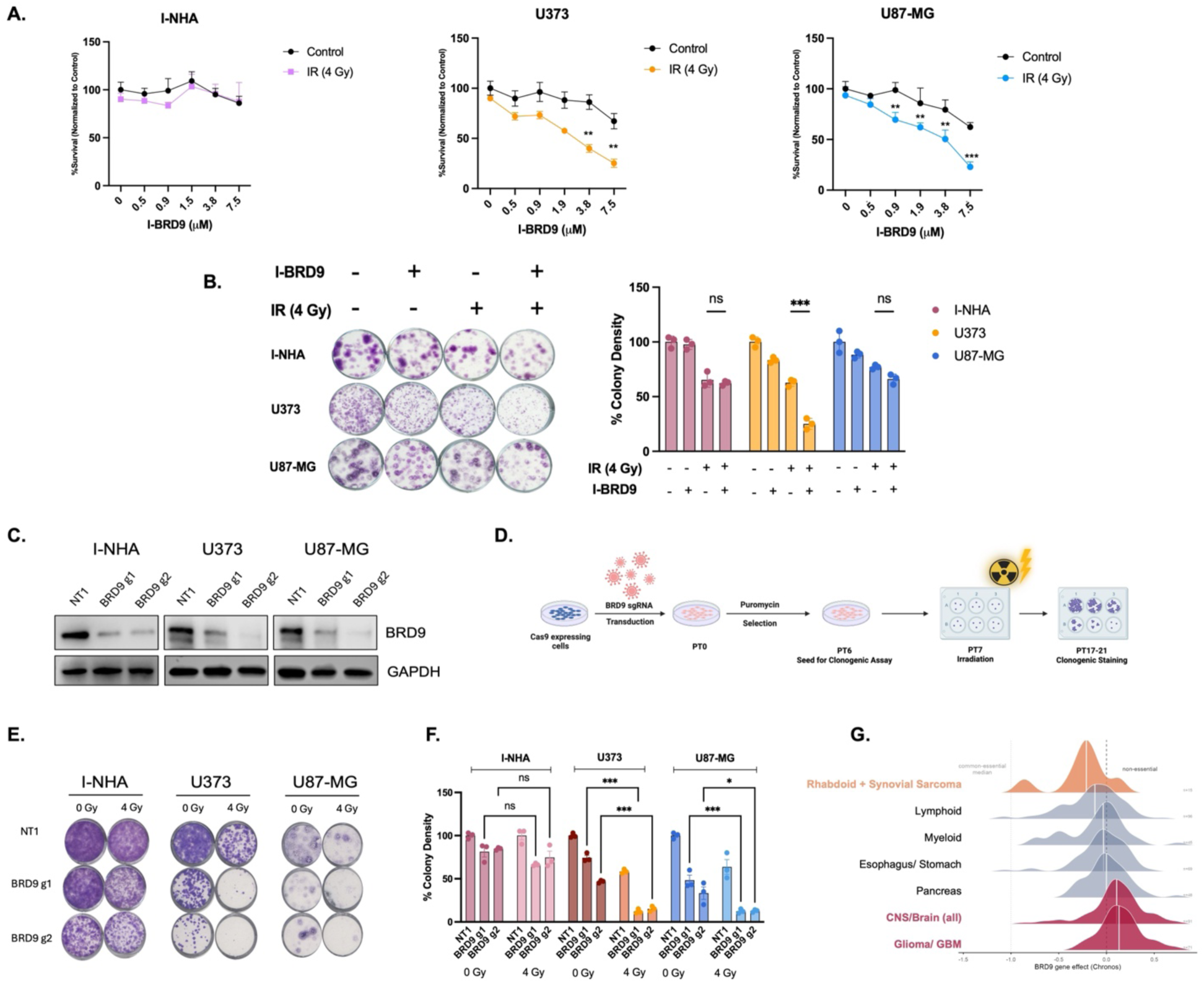
BRD9 inhibition selectively radiosensitizes glioblastoma cells but not non-malignant astrocytes. **A.** Dose–response curves showing percentage survival of I-NHA, U373, and U87-MG cells treated with I-BRD9 (0–7.5 µM) in the presence or absence of ionizing radiation (IR, 4 Gy). **B.** Representative clonogenic assay images and quantification of colony formation in I-NHA, U373, and U87-MG cells treated with I-BRD9 (2 µM) and IR (4 Gy). Data are presented as mean ± SEM**. C.** Immunoblot analysis confirming BRD9 protein depletion in I-NHA, U373, and U87-MG cells expressing a non-targeting (NT) or two independent BRD9-targeting sgRNAs. **D.** Representation of experimental workflow of clonogenic assay for BRD9 KO cells. Generated in BioRender. **E.** Representative clonogenic assay images of I-NHA, U373, and U87-MG NT and BRD9 KO cells following irradiation. **F.** Quantification of clonogenic survival from NT1 and BRD9 KO cells across all three cell lines. **G.** Distribution of BRD9 gene-effect scores (Chronos) across cancer lineages in the DepMap CRISPR screening dataset. BRD9 scores as an essential gene in rhabdoid and synovial sarcoma lines (orange) but is non-essential in CNS/brain lineages, including glioma and glioblastoma (red).

Consistent with a non-oncogene dependency rather than a classical essential gene, analysis of genome-wide CRISPR screening data from the Cancer Dependency Map (DepMap) showed that *BRD9* scores as a strong dependency in rhabdoid and synovial sarcoma lineages, tumors driven by SWI/SNF perturbation, but is largely non-essential across CNS lineages, including glioma and glioblastoma, where BRD9 gene-effect scores cluster near the non-essential threshold **(Figure 3G).** Although BRD9 is dispensable for the baseline proliferation of GBM cells, our results indicate that it becomes selectively required under the stress imposed by radiotherapy.

### BRD9 perturbation suppresses a MYC-associated transcriptional and translational program

To define the transcriptomic alterations and molecular mechanism underlying BRD9-dependent radiosensitization, we performed RNA-sequencing after pharmacological inhibition (I-BRD9, 2 µM) and genetic knockout of *BRD9* in both U373 and U87-MG cells, generating four independent transcriptomic datasets. Differential expression analysis in I-BRD9-treated U373 cells revealed downregulation of genes involved in biosynthetic and translational processes **(Figure 4A)**. Pre-ranked gene set enrichment analysis (GSEA) identified *MYC*-target signatures among the most strongly negatively enriched pathways, alongside gene sets for unfolded protein response, *MTORC1* Signalling and Epithelial and Mesenchymal Transition **(Figure 4B).** A cross-condition GSEA dot matrix confirmed that MYC-target, translation, and ribosome-biogenesis programs were coherently suppressed in both U373 and U87-MG cells **(Figure 4C).**

**Figure 4.**
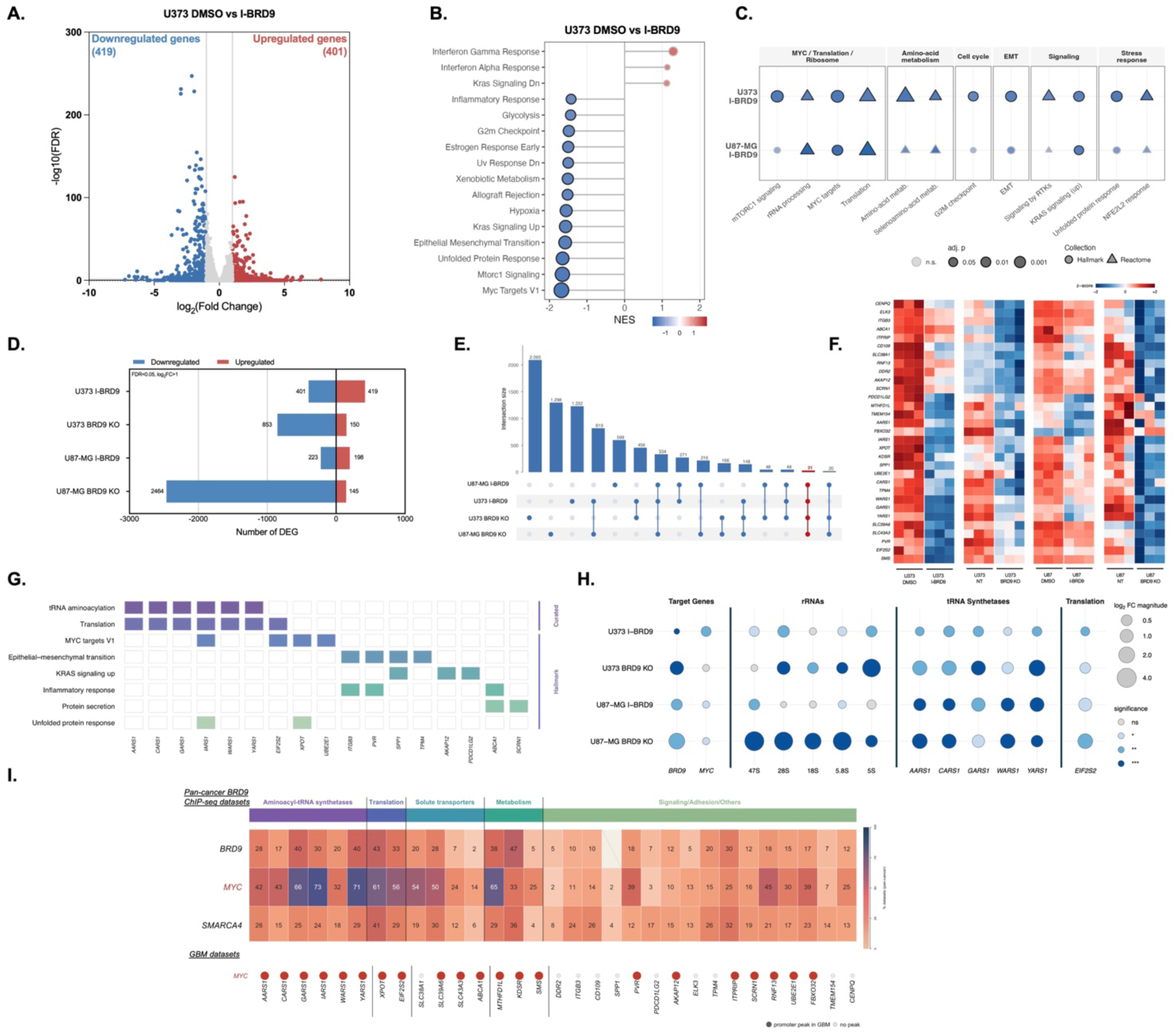
BRD9 perturbation suppresses MYC-associated transcriptional and translational programs in glioblastoma. **A.** Volcano plot showing differentially expressed genes in U373 cells following I-BRD9 treatment (2 µM, 72 h) compared to DMSO control. Significantly downregulated genes (log2FC < −1, FDR < 0.05) are shown in blue; upregulated genes (log2FC > 1, FDR < 0.05) are shown in red. **B.** GSEA dot plot highlighting enriched pathways upon BRD9 inhibition in U373 cells. Dot size represents −log10(FDR); x-axis shows normalized enrichment score (NES). **C.** Dot matrix summarizing GSEA results for U373 and U87-MG cells treated with I-BRD9 across six functional categories. Dot colour indicates NES (blue, negative; red, positive), dot size indicates adjusted p-value, and dot shape indicates the source collection (circle, Hallmark; triangle, Reactome). Faded dots denote non-significant enrichment (adjusted p ≥ 0.05). **D.** Number of significantly differentially expressed genes (FDR < 0.05, |log2FC| > 1) across the four BRD9 perturbation conditions. Downregulated genes are shown in blue and upregulated genes in red. **E.** UpSet plot of the overlap between downregulated gene sets across the four perturbation conditions. Bars show intersection sizes; connected dots below indicate which conditions contribute to each intersection. Thirty-one genes are commonly downregulated across all four conditions and define the conserved BRD9-dependent core used in subsequent analyses. **F.** Expression heatmap (z-scored across samples) of the 31 conserved BRD9-dependent core genes in U373 and U87-MG cells under control and BRD9-perturbed conditions. Rows are individual genes; columns are experimental conditions. **G.** Pathway membership tile chart for the conserved BRD9-dependent downregulated genes. Rows are curated and Hallmark pathway annotations; columns are individual genes. **H.** Expression levels of of selected MYC/BRD9 target genes, rRNA species, and tRNA synthetases and EIF2S2 across all four BRD9 perturbation conditions relative to respective controls. **I.** ChIP-Atlas analysis of transcription-factor occupancy at the promoters of the 31 conserved BRD9-dependent core genes. The upper heatmap shows the percentage of publicly deposited ChIP-seq datasets in which each factor (BRD9, MYC, SMARCA4) has a peak at each gene promoter; genes are grouped by functional category above. The lower dot panel indicates whether a promoter peak was detected for MYC in glioblastoma-derived datasets specifically.

Comparison across all four perturbation conditions demonstrated a highly consistent downregulation pattern **(Figure 4D; Supplementary Figure S5),** and intersection analysis defined a conserved core of 31 genes commonly downregulated across all four datasets **(Figure 4E,F),** indicating a BRD9-dependent program robust to perturbation modality and cell line. Pathway annotation of these 31 genes confirmed strong enrichment for tRNA aminoacylation, translation, and MYC-associated processes **(Figure 4G).** Quantitative RT-PCR validation of representative targets, rRNA species (47S, 28S, 18S, 5.8S, 5S), tRNA synthetases (*AARS1, CARS1, GARS1, WARS1, YARS1*), and the translation factor *EIF2S2* confirmed consistent downregulation across all four perturbation conditions **(Figure 4H).** To test whether the 31 genes commonly downregulated upon BRD9 inhibition and knockout are direct MYC targets, we scored promoter occupancy (peaks within ±1 kb of a TSS) in public ChIP-seq from ChIP-Atlas for BRD9 and MYC, and the positive control SMARCA4 (**Figure 4I, top)**. Across all cancer contexts, MYC, occupied all 31 core promoters, as did SMARCA4. Then restricting to glioblastoma ChIP-seq (U-87 MG, GSC316, and KNS-42), the MYC bound 21 of core promoters (**Figure 4I, bottom).** All eight tRNA-synthetase and translation genes were bound by every factor assayed in glioblastoma, whereas binding across the signaling/adhesion genes was more variable. Extending this analysis to the broader MYC network, MYC, MYCN and MAX each occupied the promoters of the majority of the 31 core genes across pan-cancer ChIP-seq datasets, and MYC/MAX promoter peaks were likewise recovered at these genes in glioblastoma specifically (**Supplementary Figure S6**). Together these data indicate that genes requiring BRD9 in glioblastoma are direct MYC targets enriched for the tRNA-synthetase and translation program, positioning BRD9 as a chromatin dependency sustaining MYC-driven anabolic output.

Collectively, these findings establish that BRD9 sustains a MYC-associated transcriptional program supporting translational capacity in GBM cells, and that disrupting this axis produces a coordinated suppression of protein-synthesis-related gene expression.

### Ectopic MYC expression rescues BRD9-inhibition mediated radiosensitization

To test whether suppression of MYC is causally responsible for BRD9-inhibition mediated radiosensitization, we ectopically overexpressed MYC in U373 cells, confirming overexpression at the mRNA and protein levels relative to wild-type (WT) and empty-vector (EV) controls (**Figure 5A,B**). MYC overexpression induced translation- and ribosome-biogenesis-associated genes, including rRNA species, tRNA synthetases, and EIF2S2, showing an expression pattern opposite to that observed upon BRD9 inhibition (**Figure 5C**), supporting a role for MYC in regulating the translational programs affected by BRD9 inhibition. Functionally, MYC overexpression shifted the I-BRD9 dose–response curve, increasing the IC50 relative to control cells (**Figure 5D**). Consistent with this, MYC overexpression substantially attenuated the loss of clonogenic survival induced by combined I-BRD9 and irradiation, with the combination producing a markedly smaller effect in MYC-OE than in control cells (**Figure 5E,F**).

**Figure 5.**
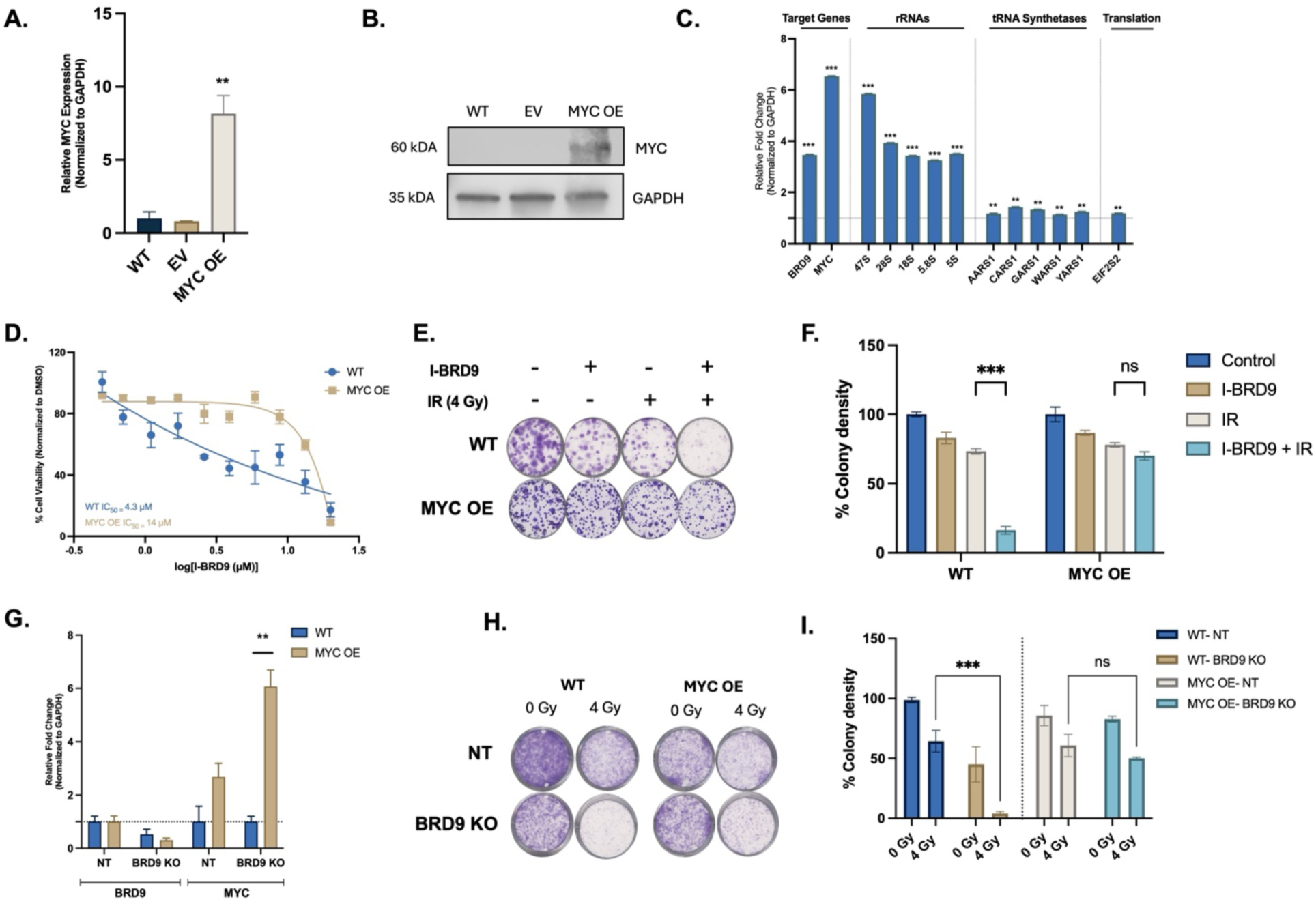
MYC overexpression rescues BRD9-inhibition mediated radiosensitization in glioblastoma cells. **A.** Relative MYC mRNA expression in U373 wild-type (WT), empty-vector (EV), and MYC-overexpressing (MYC OE) cells, measured by qRT-PCR and normalized to GAPDH. **B.**Immunoblot confirming MYC protein overexpression in U373 MYC OE cells relative to WT and EV controls. GAPDH serves as loading control. **C.** Relative mRNA expression of BRD9, MYC, rRNA species, tRNA synthetases, and EIF2S2 in MYC OE cells relative to EV controls, normalized to GAPDH. **D.** I-BRD9 dose–response curves for U373 WT and MYC OE cells. Fitted IC50 values are indicated. **E.** Representative clonogenic survival images and **(F)** quantification of colony density for U373 WT and MYC OE cells treated with I-BRD9 (2 µM) and/or 4 Gy irradiation as indicated. **G.** Relative BRD9 and MYC mRNA expression in U373 NT1 control and BRD9-g1 knockout cells, with and without MYC overexpression, measured by qRT-PCR and normalized to GAPDH. **H.** Representative clonogenic survival images and **(I)** quantification of colony density for U373 WT and MYC OE cells carrying NT1 or BRD9-g1 sgRNAs, with or without 4 Gy irradiation.

To determine whether this rescue effect extends to genetic BRD9 loss, we overexpressed MYC in U373 BRD9 KO cells (**Figure 5G**). Clonogenic assays showed that MYC overexpression markedly attenuated the reduction in post-irradiation colony formation associated with BRD9 loss (**Figure 5H,I**). Together, the pharmacological and genetic findings demonstrate that MYC overexpression reduces the radiosensitizing effects of BRD9 perturbation, supporting a functional link between BRD9, MYC and radiotherapy response in glioblastoma.

### The BRD9–MYC axis is linked to therapeutic response in glioblastoma patients and BRD9 inhibition radiosensitizes patient-derived spheroids

To evaluate the clinical relevance of the BRD9–MYC axis, we first examined whether BRD9 and MYC expression differ between primary and recurrent glioblastoma. In the TCGA GBM microarray dataset (HG-U133A platform), both BRD9 and MYC mRNA were significantly higher in recurrent than in primary tumors (BRD9, p = 0.006; MYC, p = 0.003; primary n = 497, recurrent n = 16) (**Figure 6A**). Although the recurrent sample size is limited, this pattern is directionally consistent with radiotherapy selecting for BRD9–MYC-high states. We next assessed BRD9–MYC co-expression across fourteen independent glioma and glioblastoma cohorts. Spearman correlation analysis revealed a significant positive association in nine cohorts (p < 0.05), spanning multiple platforms and tumor grades, including glioblastoma (TCGA, Cell 2013; n = 152, p < 0.001), diffuse glioma (GLASS Consortium; n = 79 and n = 355, both p < 0.001), diffuse glioma (TCGA GDC; n = 530, p < 0.05), glioblastoma multiforme (TCGA PanCancer Atlas; n = 160, p < 0.01), brain lower-grade glioma (TCGA PanCancer Atlas; n = 514, p < 0.001), CNS/brain cancer (CPTAC GDC; n = 221, p < 0.01), and brain tumor PDXs (Mayo Clinic; n = 66, p < 0.05); the remaining, generally smaller or mixed-histology cohorts showed positive but non-significant correlations (**Figure 6B, Supplementary Figure S7**). The strongest co-expression was observed in the TCGA GBM cohort (Cell 2013; Spearman ρ = 0.55, p < 0.0001) (**Figure 6C**).

**Figure 6.**
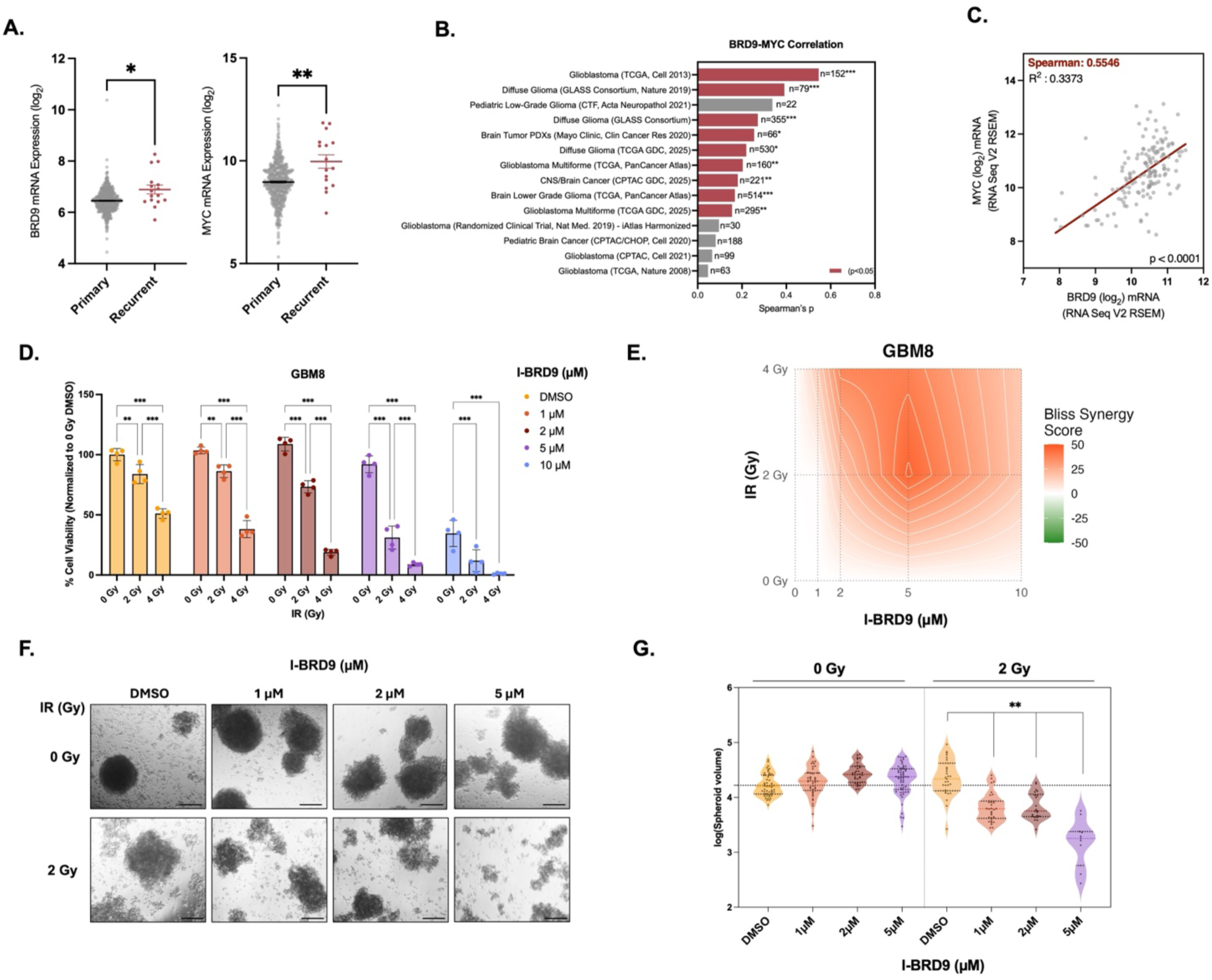
The BRD9–MYC axis is linked to therapeutic response in glioblastoma patients, and BRD9 inhibition radiosensitizes patient-derived spheroids. **A.** BRD9 and MYC mRNA expression in primary (n = 497) versus recurrent (n = 16) glioblastoma tumours from the TCGA GBM microarray dataset (HG-U133A platform). Each point represents one tumour; horizontal lines indicate mean ± SEM. Significance was assessed by two-tailed Mann–Whitney test. **B.** Spearman correlation coefficients between BRD9 and MYC mRNA expression across fourteen publicly available glioma and glioblastoma cohorts, ordered by effect size. Cohort sample sizes are indicated to the right of each bar. Red bars denote statistically significant correlations (p < 0.05); grey bars denote non-significant correlations. **C.** Scatter plot of MYC versus BRD9 mRNA expression (log2 RNA-Seq V2 RSEM) in the TCGA GBM cohort. **D.** Cell viability of GBM8 patient-derived spheroids treated with I-BRD9 (0, 1, 2, 5, or 10 µM) and ionizing radiation (0, 2, or 4 Gy), expressed as a percentage of the DMSO control and normalized to the 0 Gy condition within each drug concentration. **E.** Two-dimensional Bliss synergy landscape for the I-BRD9 × ionizing radiation combination in GBM8 spheroids, computed with SynergyFinder from the viability matrix in panel E. Red indicates synergy (Bliss score > 0) and green indicates antagonism (Bliss score < 0). **F.** Representative brightfield images of GBM8 spheroids treated with DMSO or I-BRD9 (1, 2, or 5 µM) in the absence (0 Gy) or presence (2 Gy) of ionizing radiation. Scale bar, *20* µm. **G.** Quantification of spheroid size (log-transformed) for GBM8 spheroids across the treatment conditions in panel G, at 0 Gy and 2 Gy.

To test directly whether such tumors are therapeutically vulnerable to BRD9 inhibition, we turned to a patient-derived model. In three-dimensional GBM8 spheroids, I-BRD9 reduced viability in a concentration-dependent manner, and this effect was markedly potentiated by ionizing radiation across the tested dose range (**Figure 6D**). Two-dimensional Bliss synergy analysis confirmed that I-BRD9 treatment and radiation interact synergistically in this patient-derived context (**Figure 6E**). Irradiation alone had only a limited effect on spheroid viability but partially disrupted spheroid architecture, yielding a looser, less compact structure; combined I-BRD9 and irradiation caused pronounced spheroid disaggregation on brightfield imaging (**Figure 6F**). To quantify these effects, spheroid volume was calculated from brightfield images on day 10, and the greatest reduction in spheroid volume occurred with the combination of increasing I-BRD9 concentration and irradiation (**Figure 6G**). These results indicate that BRD9 inhibition radiosensitizes a patient-derived glioblastoma spheroid model, consistent with the effects observed in established cell lines and supporting further evaluation of BRD9 inhibition as a radiosensitizing strategy in glioblastoma.

## DISCUSSION

Our study identifies BRD9 inhibition as a priming radiosensitization strategy in glioblastoma that is mechanistically grounded in the suppression of MYC-dependent translational programs required for post-irradiation survival. Several features distinguish this finding from conventional epigenetic radiosensitization approaches. First, the vulnerability was identified through a focused screen conducted specifically under irradiation, selecting for context-dependent dependencies rather than baseline cytotoxicity. Second, BRD9 inhibition exerts minimal single-agent toxicity yet selectively potentiates irradiation-induced apoptosis in malignant cells while sparing non-malignant astrocytes, and the drug–radiation interaction is synergistic by Bliss analysis across three structurally distinct BRD9 inhibitors. Third, pre-treatment is markedly more effective than co- or post-treatment, indicating that BRD9 inhibition might establish a transcriptionally vulnerable state before irradiation, consistent with the non-oncogene addiction framework, in which cancer cells depend on non-mutated stress-support genes that become selectively essential under therapeutic pressure^39,40^. The DepMap analysis reinforces this framing: BRD9 is a strong dependency in SWI/SNF-driven sarcomas but non-essential across CNS lineages, underscoring that its requirement in glioblastoma is conditional on radiation stress rather than constitutive.

Transcriptomic profiling across four independent perturbation conditions converges on a coherent mechanism: BRD9 sustains a MYC-associated program that maintains expression of translation-associated genes, spanning ribosome biogenesis, rRNA processing, tRNA aminoacylation, and translation initiation. A conserved 31-gene core, downregulated across all conditions and enriched for these pathways, is occupied at the chromatin level by MYC and the ncBAF ATPase SMARCA4 in neural-lineage ChIP-seq datasets. Functionally, ectopic MYC overexpression restores the translational program and rescues BRD9-dependent radiosensitization both pharmacologically and after genetic BRD9 loss, supporting the sufficiency of MYC rather than proving its necessity. This BRD9–MYC–translation axis mirrors findings in acute myeloid leukemia and multiple myeloma, where BRD9 sustains MYC output at superenhancers and occupies ribosome-biogenesis promoters in a complex with BRD4 and MYC^41,42^, suggesting that BRD9 is a broadly utilized chromatin factor linking H3K27ac-marked enhancer state to MYC-driven output. This biology aligns with the broader concept of translational addiction in MYC-activated cancers, in which coordinated upregulation of ribosome biogenesis, tRNA charging, and translation initiation sustains oncogenic protein output^43,44^.

Regarding the DNA-damage phenotype, the sustained γH2AX and 53BP1 foci at 24 hours after combination treatment most likely reflect impaired biosynthetic capacity for the stress response rather than a primary DNA-repair defect. The strongest phenotype is enhanced apoptosis, in which BRD9 inhibition reduces translational rather than repair gene programs. The required priming timing argues against direct repair inhibition, since post-treatment would otherwise be equally effective. Our model remains consistent with prior reports that BRD9 can directly facilitate RAD51–RAD54 complex assembly in some non-glial tumors^27^; under our conditions, however, baseline repair and early γH2AX clearance were not substantially altered by BRD9 inhibition alone. The kinetic phenotype emerged specifically in the post-irradiation recovery window when translational support of stress-response proteins is most demanding. The coordinated downregulation of tRNA aminoacylation enzymes alongside ribosomal and rRNA-processing genes further raises the possibility that BRD9 perturbation engages or unmasks the integrated stress response, and whether stress signaling cooperates with BRD9 inhibition to amplify post-irradiation apoptosis is an attractive hypothesis for direct testing.

The clinical coherence of the BRD9–MYC axis is supported by patient data: BRD9 and MYC are elevated at recurrence and co-expressed across nine of fourteen independent cohorts examined, indicating that the axis is engaged in a substantial subset of tumours. Importantly, we complement these correlative observations with functional patient-derived material: in GBM8 spheroids, BRD9 inhibition synergized with irradiation to reduce viability and spheroid volume, providing patient-derived support for the combination beyond established cell lines. In glioblastoma more broadly, MYC-family aberrations occur in defined tumor subsets and have been associated with aggressive behavior and limited therapeutic options^45^, consistent with a rationale for targeting MYC-supporting dependencies such as BRD9 in this disease.

There are several limitations in our study. Our functional data are derived from established cell lines and a single patient-derived spheroid model; additional patient-derived organoids or in vivo experiments are needed to assess blood-brain-barrier penetration, tolerability, and translational feasibility. We did not perform BRD9-specific ChIP-seq in our cell models and therefore interpret BRD9 as supporting MYC-dependent programs through ncBAF rather than asserting direct BRD9 binding at these promoters. All radiosensitization experiments used single-dose irradiation without concurrent temozolomide, so whether the synergy persists under fractionated radiotherapy and concurrent temozolomide, the components of standard-of-care chemoradiation, remains to be tested. Our rescue experiments establish that restoring MYC is sufficient to reverse radiosensitization but do not test necessity, and parallel ncBAF-, integrated-stress-response-, and RAD51–RAD54-dependent contributions were not formally excluded. It is notable that BET inhibitors targeting BRD4 have shown disappointing single-agent activity in glioblastoma, partly due to rapid adaptive resistance^46^. The BRD9–ncBAF axis defined here is mechanistically distinct and shows limited effects as a single agent target, while BRD9 inhibition enhances radiation response through a MYC-associated translational program, suggesting that combining BRD9-targeted therapy with radiotherapy may offer an alternative strategy to BET-inhibitor monotherapy.

The emergence of clinical-grade BRD9 degraders adds translational weight to these findings. Heterobifunctional PROTAC degraders such as FHD-609 (Foghorn Therapeutics) and the orally bioavailable CFT8634 (C4 Therapeutics) achieve potent and selective BRD9 degradation and have advanced into first-in-human trials in synovial sarcoma and SMARCB1-deficient tumors ^47^. To date, however, single-agent clinical activity has been modest, and the FHD-609 program encountered dose-limiting prolongation, underscoring that BRD9 inhibition alone may be insufficient in tumors that do not constitutively depend on BRD9 ^48^. This clinical experience is consistent with our observation that BRD9 is largely non-essential at baseline and becomes a vulnerability only under therapeutic stress: rather than a monotherapy, BRD9 degradation may be most valuable as a sensitizing partner that exposes a conditional, irradiation-induced context. In this case, pairing a BRD9 degrader with radiotherapy, rather than utilizing it as a stand-alone agent, offers a mechanistically rational path to exploit the priming effect we describe. Realizing this in glioblastoma will additionally require BRD9 degraders with demonstrated central nervous system penetration, as the blood–brain barrier remains a key hurdle for the current generation of these compounds and an important direction for future medicinal-chemistry efforts^49–52^.In summary, BRD9 inhibition disrupts a MYC-associated gene-expression program and enhances the response of glioblastoma cells to irradiation, while having comparatively limited effects on non-malignant cells. The consistent radiosensitizing effects observed across multiple BRD9 inhibitors, together with the ability of MYC overexpression to attenuate these effects, support a functional link between BRD9, MYC, and radiation response. These findings provide a rationale for further preclinical evaluation of BRD9 inhibition in combination with radiotherapy (**Figure 7**).

**Figure 7.**
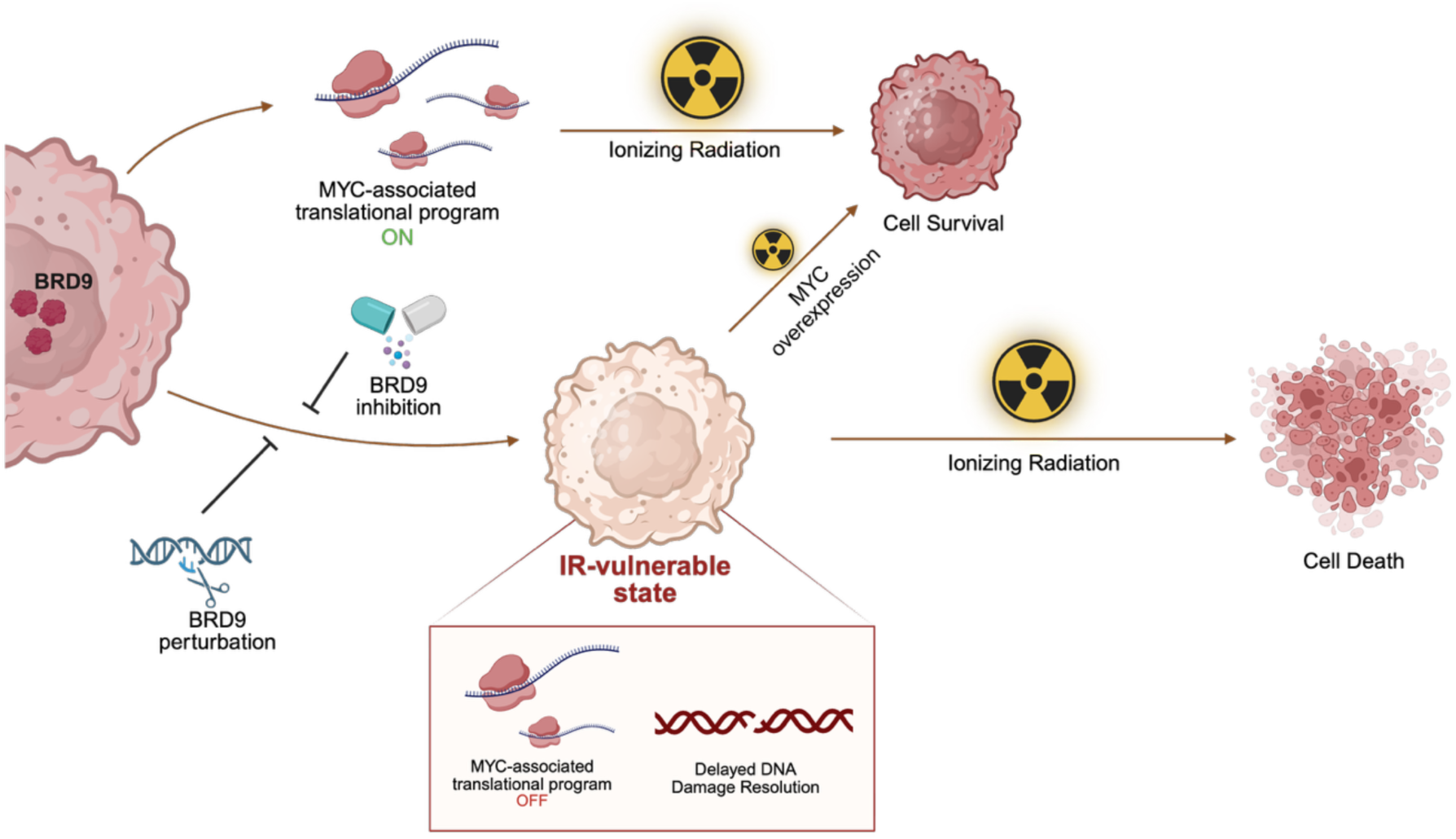
Graphical Abstract. In glioblastoma cells, BRD9 activity sustains MYC-associated translational programs that enable survival following irradiation-induced stress, thereby contributing to radiotherapy resistance. BRD9 inhibition collapses these MYC-dependent translational programs, delays the resolution of radiation-induced DNA damage, and increases apoptotic cell death, collectively enhancing the efficacy of ionizing radiation. Ectopic MYC overexpression partially increases translational gene expression and attenuates BRD9 inhibition-mediated radiosensitization, establishing MYC as a central functional mediator of this axis.

## Data Availability

RNA sequencing data is deposited to the EMBL Array Express database with the accession number E-MTAB-17302. The following figures were created using BioRender.com and licensed for publication (Fig.1B: JF29KB8EHE, Fig.1K: QQ29KB8JD5, Fig.2D: GE29ZW1LY1, Fig.3E: AV2A0Q0852, Fig.S1D: SV29KBA9D0, Fig. 7: QH2A1APUQE)

## Author Contributions

Study design: T.B.O. and N.D.; Methodology: T.B.O, U.S., and N.D.; data generation, resources and analysis: N.D, S.A.Ç, B.N.K, U.O.; data interpretation: T.B.O., N.D., S.A.Ç, B.N.K, U.S. and A.P.C. drafted the manuscript: T.B.O., N.D.; approved final manuscript: all authors.

## Acknowledgements

The authors gratefully acknowledge the use of the services and facilities of the Koç University Research Center for Translational Medicine (KUTTAM), funded by the Presidency of Turkey, Head of Strategy and Budget. Authors thank Neelam Mehta and Altar Özbıyık for sequencing and irradiation experiments. The work was supported through the LEAN (Leducq Epigenetics Atherosclerosis Network) program grant of the Leducq Foundation (UO), Cancer Research UK (U.O.), the Bone Cancer Research Trust (A.P.C. and U.O.), the Chan Zuckerberg Initiative (A.P.C.), and the MRC Career Development Fellowship MR/V010182/1 (A.P.C.). Authors thank Koç University Animal Research Facility (KUARF) and Koç University Hospital Radiation Oncology Department for irradiation experiments. This study is funded by International Centre for Genetic Engineering and Biotechnology (ICGEB) and Eczacıbaşı Scientific Research Grant.

## Conflict of Interest

U.O. and A.P.C. are co-founders of Entelo Bio. All other authors declare no competing interests.

## Notes

### Competing Interest Statement

The authors have declared no competing interest.

