## Supplementary Materials for "BRD9 inhibition induces selective radiosensitivity in glioblastoma through MYC pathway modulation"

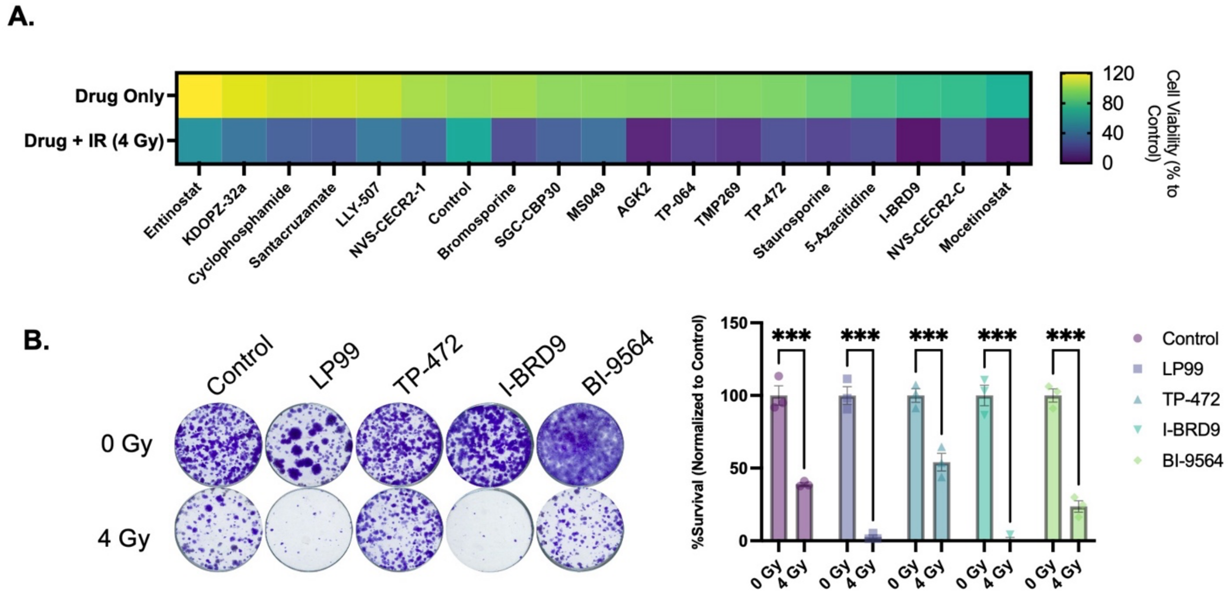


**Supplementary Figure 1. Validation of BRD9 radiosensitization using additional BRD9-targeting compounds.**

**A.** Heatmap of the top 20 radiosensitizing compounds ranked by their ability to preferentially reduce cell viability in U373 cells in combination with ionizing radiation (IR, 4 Gy) relative to drug treatment alone. Cell viability is expressed as percentage relative to untreated controls. BRD9-targeting compounds are highlighted. **B.** Representative clonogenic survival images (left) and colony-density quantification (right) for U373 cells treated with LP99, TP-472, I-BRD9 or BI-9564 in the absence or presence of ionizing IR. Colony density was normalized to DMSO-treated, non-irradiated controls.

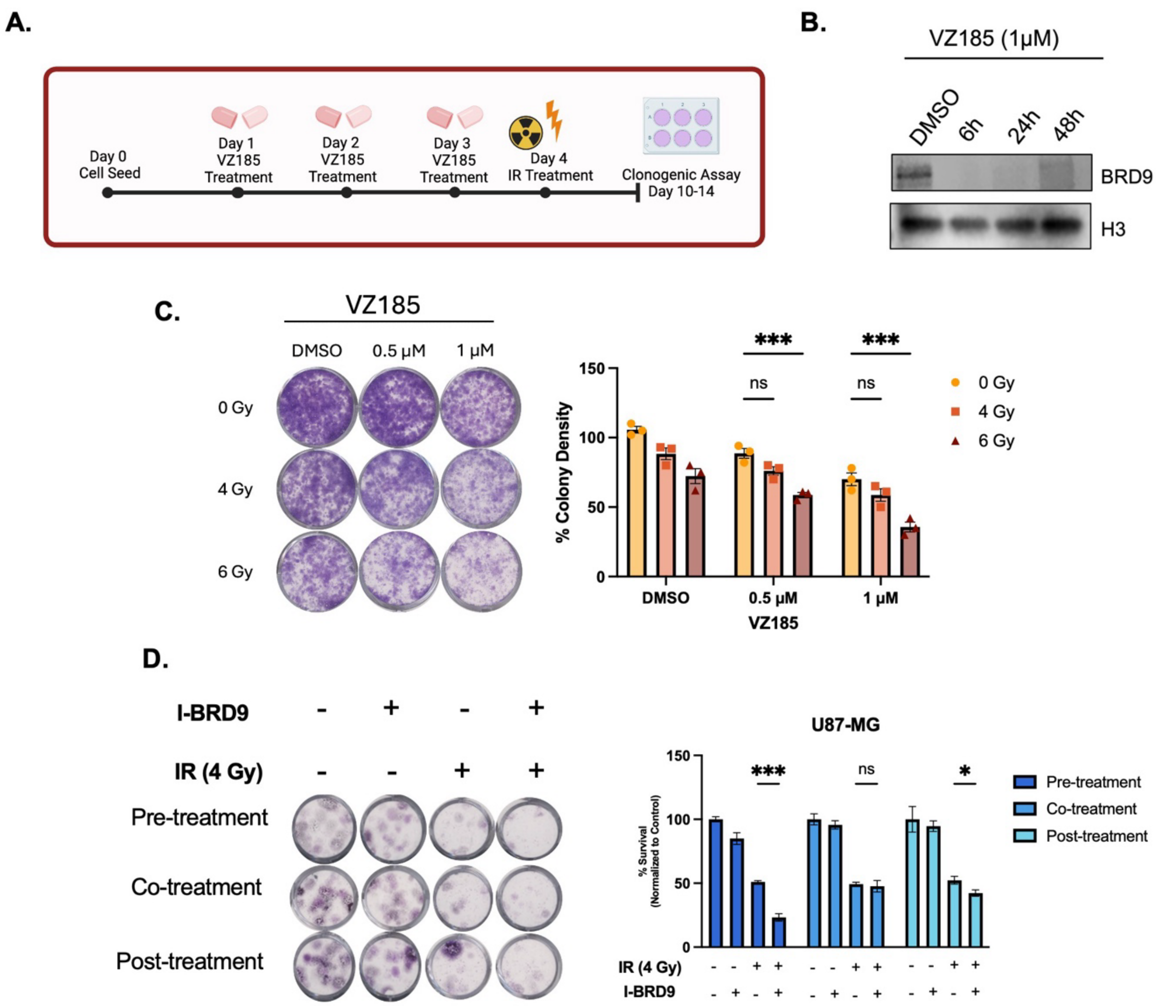


**Supplementary Figure S2. BRD9 degradation and treatment-timing dependence of BRD9-inhibition mediated radiosensitization in glioblastoma cells**

**A.** Schematic overview of the treatment schedule used for evaluation of the BRD7/BRD9 PROTAC degrader VZ185. Cells were pre-treated with VZ185 for 72 hours prior to irradiation. Generated in BioRender. **B.** Immunoblot analysis confirming BRD9 protein depletion in U373 cells following treatment with VZ185 at the indicated concentrations. **C.** Representative clonogenic assay images and corresponding quantification of U373 cells treated with VZ185 and exposed to increasing doses of ionizing radiation (0–6 Gy). **D.** Representative clonogenic survival images and quantification of normalized colony density for U87-MG cells treated according to each schedule with I-BRD9 (2 μM) and/or 4 Gy irradiation, as indicated below the axis.

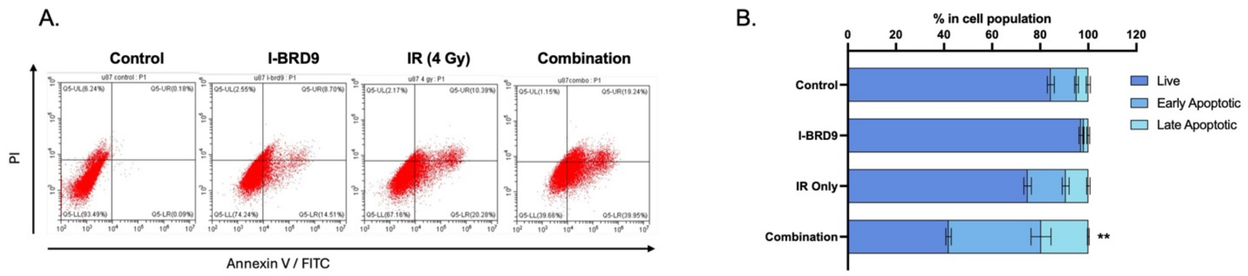


**Supplementary Figure S3. Validation of apoptosis induction by BRD9 inhibition and irradiation in U87-MG cells.**

**A.** Representative Annexin V/PI flow cytometry plots of U87-MG cells treated with vehicle control (DMSO), I-BRD9 (2 μM), ionizing radiation (IR, 4 Gy), or the combination of I-BRD9 and IR. **B.** Quantification of live, early apoptotic, and late apoptotic cell populations from panel A following the indicated treatments. Data are presented as mean ± SEM.

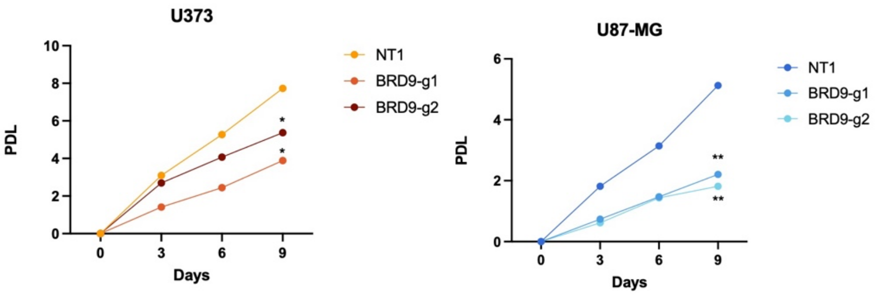


**Supplementary Figure S4. Effects of BRD9 loss on baseline proliferation of GBM cells.** Population doubling level (PDL) growth curves of U373 (left) and U87-MG (right) NT1 control cells and BRD9 knockout cells (BRD9-g1 and BRD9-g2) cultured under standard growth conditions without irradiation.

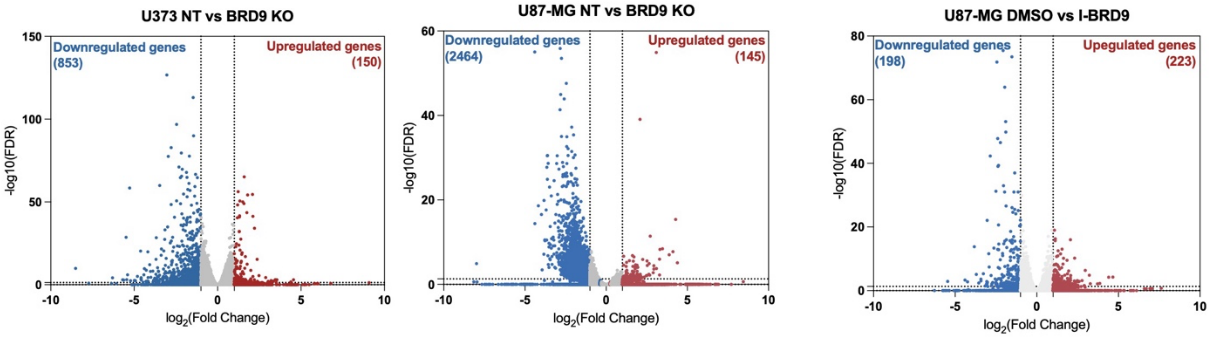


**Supplementary Figure S5. Volcano plots of differentially expressed genes across additional BRD9 perturbation conditions.**

Volcano plots showing differentially expressed genes in: U373 BRD9 knockout (BRD9 KO) vs. NT1 control cells (left); U87-MG BRD9 KO vs. NT1 control cells (center); and U87-MG cells treated with I-BRD9 (5 μM) vs. DMSO (right). Significantly downregulated genes (log2FC < −1, FDR < 0.05) are shown in blue; upregulated genes (log2FC > 1, FDR < 0.05) in red. The total number of significantly differentially expressed genes in each direction is indicated on each plot.

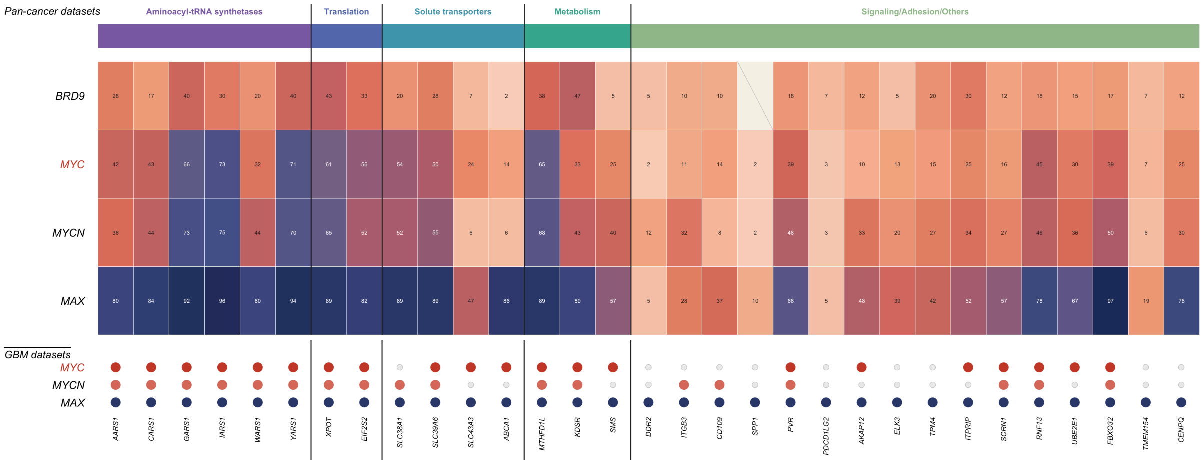


**Supplementary Figure S6. MYC-family transcription factors occupy the promoters of the BRD9-repressed 31-gene core across cancers and in glioblastoma.** Promoter occupancy of BRD9 and the MYC-family transcription factors MYC, MYCN and their obligate dimerization partner MAX across the 31 commonly downregulated core genes, grouped by functional category. *Top,* heat map showing, for each factor–gene pair, the percentage of pan-cancer ChIP-seq datasets in ChIP-Atlas (hg38, "Target Genes") in which a peak was called in the promoter-proximal region (±1 kb of the transcription start site); darker shading indicates promoter binding in a larger fraction of datasets. *Bottom,* dot plot indicating whether a promoter peak for MYC, MYCN or MAX was detected in glioblastoma ChIP-seq datasets specifically (filled circle, peak present; open circle, no peak).

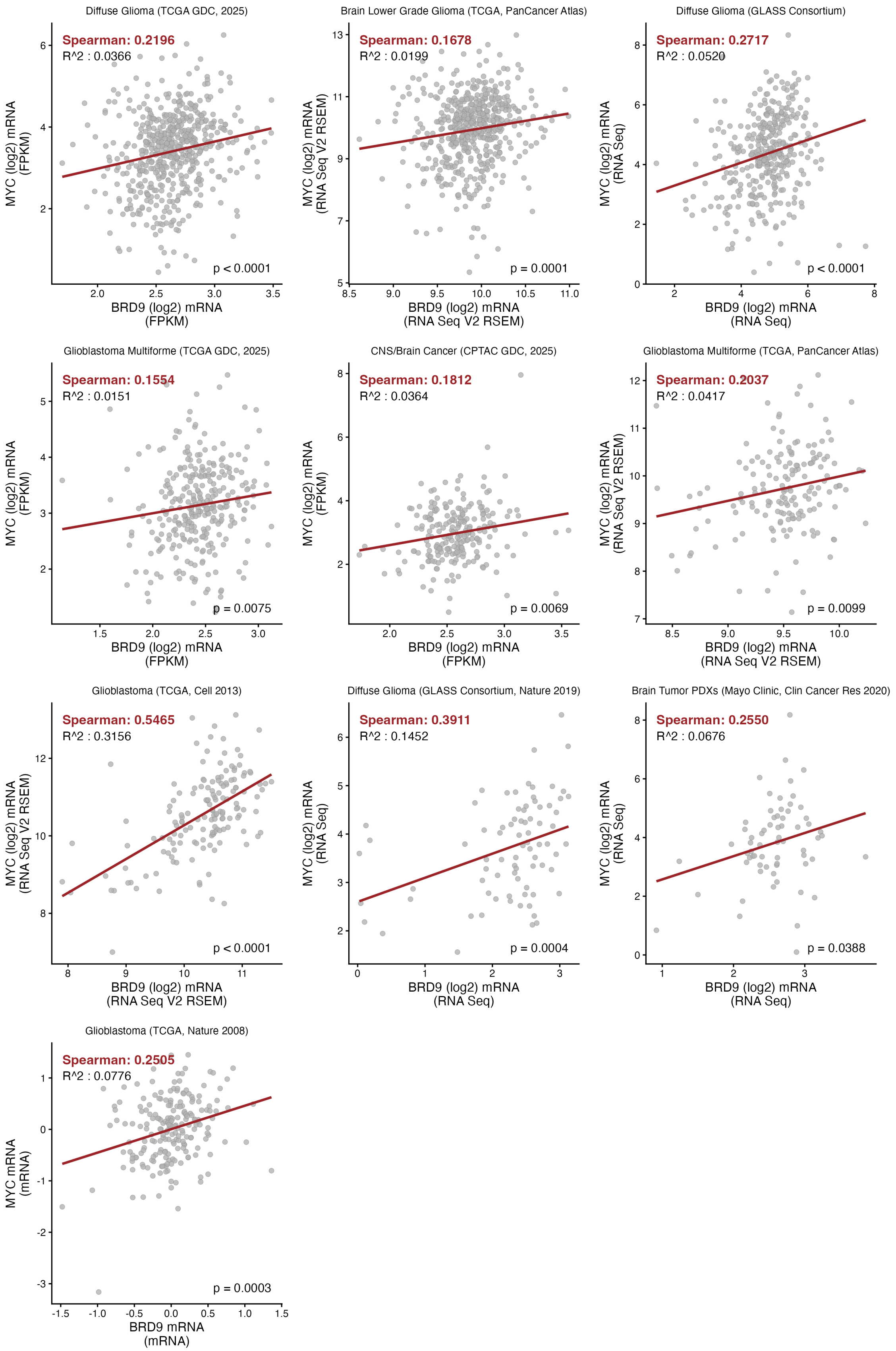
Diffuse Glioma (TCGA GDC, 2025)
Spearman: 0.2196
R^2 : 0.0366
p < 0.0001
MYC (log2) mRNA (FPKM)
BRD9 (log2) mRNA (FPKM)
Brain Lower Grade Glioma (TCGA, PanCancer Atlas)
Spearman: 0.1678
R^2 : 0.0199
p = 0.0001
MYC (log2) mRNA (RNA Seq V2 RSEM)
BRD9 (log2) mRNA (RNA Seq V2 RSEM)
Diffuse Glioma (GLASS Consortium)
Spearman: 0.2717
R^2 : 0.0520
p < 0.0001
MYC (log2) mRNA (RNA Seq)
BRD9 (log2) mRNA (RNA Seq)
Glioblastoma Multiforme (TCGA GDC, 2025)
Spearman: 0.1554
R^2 : 0.0151
p = 0.0075
MYC (log2) mRNA (FPKM)
BRD9 (log2) mRNA (FPKM)
CNS/Brain Cancer (CPTAC GDC, 2025)
Spearman: 0.1812
R^2 : 0.0364
p = 0.0069
MYC (log2) mRNA (FPKM)
BRD9 (log2) mRNA (FPKM)
Glioblastoma Multiforme (TCGA, PanCancer Atlas)
Spearman: 0.2037
R^2 : 0.0417
p = 0.0099
MYC (log2) mRNA (RNA Seq V2 RSEM)
BRD9 (log2) mRNA (RNA Seq V2 RSEM)
Glioblastoma (TCGA, Cell 2013)
Spearman: 0.5465
R^2 : 0.3156
p < 0.0001
MYC (log2) mRNA (RNA Seq V2 RSEM)
BRD9 (log2) mRNA (RNA Seq V2 RSEM)
Diffuse Glioma (GLASS Consortium, Nature 2019)
Spearman: 0.3911
R^2 : 0.1452
p = 0.0004
MYC (log2) mRNA (RNA Seq)
BRD9 (log2) mRNA (RNA Seq)
Brain Tumor PDXs (Mayo Clinic, Clin Cancer Res 2020)
Spearman: 0.2550
R^2 : 0.0676
p = 0.0388
MYC (log2) mRNA (RNA Seq)
BRD9 (log2) mRNA (RNA Seq)
Glioblastoma (TCGA, Nature 2008)
Spearman: 0.2505
R^2 : 0.0776
p = 0.0003
MYC mRNA (mRNA)
BRD9 mRNA (mRNA)

**Supplementary Figure S6. BRD9 and MYC mRNA are positively correlated across glioma patient cohorts.**

*MYC* versus *BRD9* mRNA ($\log_2$) in glioma datasets from cBioPortal; each point is one tumour, dark-red line is the linear fit. Spearman's ρ, $R^2$, and p are shown per panel. Only cohorts with a significant positive correlation (Spearman $p < 0.05$) are displayed.

**Supplementary Table 1**. Composition of chemical drug library

| No | Compounds | Class/Target | Working Con. (μM) |
|---|---|---|---|
| **1** | AMI-1 | Arginine methyltransferase - PRMT | 50 |
| **2** | SGC707 | Arginine methyltransferase - PRMT3 | 1 |
| **3** | TP-064 | Arginine methyltransferase - PRMT4 | 1 |
| **4** | TP-064N | Arginine methyltransferase - PRMT4 | 1 |
| **5** | MS049 | Arginine methyltransferase - PRMT4, PRMT6 | 1 |
| **6** | MS409N | Arginine methyltransferase - PRMT4, PRMT6 | 1 |
| **7** | GSK591 | Arginine methyltransferase - PRMT5 | 1 |
| **8** | LLY-283 | Arginine methyltransferase - PRMT5 | 1 |
| **9** | MS023 | Arginine methyltransferase - Type I PRMTs | 1 |
| **10** | GSK8814 | Bromodomains - ATAD2 | 10 |
| **11** | GSK8815 | Bromodomains - ATAD2 | 10 |
| **12** | GSK2801 | Bromodomains - BAZ2A, BAZ2B | 1 |
| **13** | BAZ2-ICR | Bromodomains - BAZ2A, BAZ2B | 1 |
| **14** | BAY-299 | Bromodomains - BRD1, TAF1 | 1 |
| **15** | RVX-208 | Bromodomains - BRD2, BRD3, BRD4, BRDT (BET, BD2) | 5 |
| **16** | (+)-JQ1 | Bromodomains - BRD2, BRD3, BRD4, BRDT (BET) | 1 |
| **17** | PFI-1 | Bromodomains - BRD2, BRD3, BRD4, BRDT (BET) | 5 |
| **18** | I-BET | Bromodomains - BRD2/3/4 | 1 |
| **19** | I-BRD9 | Bromodomains - BRD9 | 10 |
| **20** | TP-472 | Bromodomains - BRD9 | 1 |
| **21** | TP-472N | Bromodomains - BRD9 | 1 |
| **22** | LP99 | Bromodomains - BRD9, BRD7 | 1 |
| **23** | BI-9564 | Bromodomains - BRD9, BRD7 | 1 |
| **24** | GSK6853 | Bromodomains - BRPF1/2/3 | 1 |
| **25** | GSK9311 | Bromodomains - BRPF1/2/3 | 1 |
| **26** | PFI-4 | Bromodomains - BRPF1B | 1 |
| **27** | CBP/BRD4 (0383) | Bromodomains - CBP, BRD4(1) | 5 |
| **28** | NVS-CECR2-1 | Bromodomains - CECR2 | 1 |
| **29** | NVS-CECR2-C | Bromodomains - CECR2 | 1 |
| **30** | I-CBP112 | Bromodomains - CREBBP, EP300 | 1 |
| **31** | SGC-CBP30 | Bromodomains - CREBBP, EP300 | 1 |
| **32** | (-)-JQ1 (inactive) | Bromodomains - Negative control | 1 |
| **33** | Bromosporine | Bromodomains - pan-Bromodomain | 1 |
| **34** | OF-1 | Bromodomains - pan-BRPF | 5 |
| **35** | NI-57 | Bromodomains - pan-BRPF | 1 |
| **36** | GSK4027 | Bromodomains - PCAF, GCN5 | 1 |
| **37** | GSK4028 | Bromodomains - PCAF, GCN5 | 1 |
| **38** | L-Moses | Bromodomains - PCAF, GCN5 | 1 |
| **39** | D-Moses | Bromodomains - PCAF, GCN5 | 1 |
| **40** | SMARCA | Bromodomains - SMARCA, PB1 | 2,5 |
| **41** | PB1/SMARCA | Bromodomains - SMARCA, PB1 | 1 |
| **42** | PFI-3 | Bromodomains - SMARCA2/4, PB1(5) | 1 |
| **43** | TRIM24/BRPF | Bromodomains - TRIM24/BRPF | 10 |
| **44** | GSK864 | Dehydrogenase | 5 |
| **45** | 5-Azacitidine | DNA methyltransferase (DNMT) - | 10 |
| **46** | 5-Azadeoxycitidine | DNA methyltransferase (DNMT) - DNMT1/3 | 5 |
| **47** | CXD101 | HDAC - | 1 |
| **48** | PCI-24781 | HDAC - | 10 |
| **49** | Romidepsin | HDAC - | 1 |
| **50** | Mocetinostat | HDAC - | 10 |
| **51** | CI-994 | HDAC - 1,2,3,(8) | 1 |
| **52** | Valproic acid | HDAC - aliphatic acid compounds | 1000 |
| **53** | RGFP966 | HDAC - HDAC3 | 10 |
| **54** | Rocilinostat | HDAC - HDAC6 | 10 |

| | | | |
|---|---|---|---|
| **55** | Tubastatin A HCl | HDAC - HDAC6 | 10 |
| **56** | PCI-34051 | HDAC - HDAC8 | 5 |
| **57** | Belinostat | HDAC - hydroxamic acids | 5 |
| **58** | SAHA | HDAC - hydroxamic acids | 2,5 |
| **59** | Trichostatin A | HDAC - hydroxamic acids - Class I & II | 0,5 |
| **60** | Entinostat | HDAC - ortho-amino anilides | 0,5 |
| **61** | EX 527 | HDAC - SIRT1 | 1 |
| **62** | SRT1720 | HDAC - SIRT1 (indirect?) activator | 1 |
| **63** | AGK2 | HDAC - SIRT2 | 10 |
| **64** | TMP269 | HDAC -4, 5, 7 &9 | 10 |
| **65** | TMP195 | HDAC -4,5,7,9 | 1 |
| **66** | Santacruzamate | HDAC 2 | 50 |
| **67** | C646 | HAT- p300/CBP | 1 |
| **68** | A-485 | HAT- p300/CBP | 1 |
| **69** | A-486 | HAT- p300/CBP | 1 |
| **70** | Methylstat (Ester) | Histone demethylase | 2,5 |
| **71** | KDOBA67 | Histone demethylase | 10 |
| **72** | KDM5-C70 | Histone demethylase - JARID1 | 10 |
| **73** | ML324 | Histone demethylase - JMJD2E | 5 |
| **74** | (E)-JIB-04 | Histone demethylase - Pan JmjC | 0,05 |
| **75** | SGC0946 | Histone methyltransferase - DOT1L | 7,5 |
| **76** | GSK343 | Histone methyltransferase - EZH2 | 3 |
| **77** | UNC1999 | Histone methyltransferase - EZH2 | 1 |
| **78** | UNC2400 | Histone methyltransferase - EZH2 | 1 |
| **79** | CPI-360 | Histone methyltransferase – EZH1/2 | 10 |
| **80** | CPI-169 | Histone methyltransferase – EZH1/2 | 10 |
| **81** | UNC0642 | Histone methyltransferase - G9a, GLP | 1 |
| **82** | UNC0638 | Histone methyltransferase - G9a, GLP | 1 |
| **83** | A-366 | Histone methyltransferase - G9a, GLP | 2 |
| **84** | PFI-2 | Histone methyltransferase - SETD7 | 2 |
| **85** | LLY-507 | Histone methyltransferase - SMYD2 | 1 |
| **86** | BAY-598 | Histone methyltransferase - SMYD2 | 1 |
| **87** | PFI-5 | Histone methyltransferase - SMYD2 | 1 |
| **88** | Chaetocin | Histone methyltransferase - SUV39H1 | 0,05 |
| **89** | A-196 | Histone methyltransferase-SUV420H1/2 | 1 |
| **90** | K00135 | Kinase inhibitor - ATP competitive - PIM | 1 |
| **91** | 5-Iodotubercidin | Kinase inhibitor - ATP mimetic - Haspin | 1 |
| **92** | SGI-1776 | Kinase inhibitor - Haspin | 10 |
| **93** | CHR-6494 | Kinase inhibitor - Haspin | 1 |
| **94** | KDOAM-25a | Lysine demethylases - JARID | 1 |
| **95** | KDOAM32 | Lysine demethylases - JARID | 1 |
| **96** | GSK J4 | Lysine demethylases - JMJD3, UTX, JARID | 10 |
| **97** | KDOPZ-32a | Lysine demethylases - KDM5 | 1 |

| | | | |
|---|---|---|---|
| **98** | Tranylcypromine | Lysine demethylases - LSD1 | 20 |
| **99** | GSK-LSD1 | Lysine demethylases - LSD1 | 0,5 |
| **100** | GSK2879552 | Lysine demethylases - LSD1 | 10 |
| **101** | GSK J5 (inactive) | Lysine demethylases - Negative control | 10 |
| **102** | IOX1 (5-carboxy-8HQ) | Lysine demethylases - pan-2-OG | 40 |
| **103** | KDOOA012000 | Lysine demethylases KDM2 | 1 |
| **104** | A-395 | Methyl Lysine Binder - EED | 1 |
| **105** | A-395N | Methyl Lysine Binder - EED | 1 |
| **106** | UNC1215 | Methyl Lysine Binder - L3MBTL3 | 5 |
| **107** | OICR-9429 | Methyl Lysine Binder - WDR5 | 1 |
| **108** | TDO20824a | Methyl Lysine Binder/tudor domain | 1 |
| **109** | TDO20826a | Methyl Lysine Binder/tudor domain | 1 |
| **110** | GSK484 | Peptidyl arginine deiminase (PAD4) | 1 |
| **111** | GSK106 | Peptidyl arginine deiminase (PAD4) | 1 |
| **112** | Olaparib | Poly ADP ribose polymerase (PARP) | 1 |
| **113** | Rucaparib | Poly ADP ribose polymerase (PARP) | 10 |
| **114** | IOX2 | Prolyl-Hydroxylases - PHD2 (EGLN1) | 10 |
| **115** | MAZ1805 | tRNA Synthetase | 1 |
| **116** | MAZ1392 | tRNA Synthetase | 1 |
| **117** | Bortezomib | Proteasome | 0,1 |
| **118** | Carfilzomib | Proteasome | 0,1 |
| **119** | Temozolomide | | 50 |
| **120** | Doxorubicin | Topoisomerase 2 | 1 |
| **121** | Taxol | Tubulin | 0,01 |
| **122** | Vincristine | Tubulin | 0,01 |
| **123** | Staurosporine | Protein Kinases | 0,01 |
| **124** | Cyclophosphamide | DNA Crosslink | 10 |
| **125** | MS-275 | HDAC | 2,5 |

**Supplementary Table 2**. Antibodies used in this study

| Antibody | Brand/Catalog number |
|---|---|
| **GAPDH** | Abcam/ab9485 |
| **BRD9** | Invitrogen/PA5-110872 |
| **MYC (D3N8F)** | Cell Signaling/13987 |
| **Phospho-Histone H2A.X (Ser139)** | Millipore/05-636 |
| **53BP1** | Novus/NB100-304 |
| **Alexa Fluor® 488 goat Anti-mouse IgG** | Invitrogen/A11029 |
| **Alexa Fluor® 594 goat Anti-rabbit IgG** | Invitrogen/A11058 |
| **Histone H3** | Cell Signalling/4499S |

### **Supplementary Table 3**. qPCR primers used in this study

| Gene | Forward Primer (5' to 3') | Reverse Primer (5' to 3') |
|---|---|---|
| **BRD9** | ATGTTCCATGAAGCCTCCAG | AGCTCCTTCTTCACCTTCCC |
| **MYC** | TACAACACCCGAGCAAGGAC | GAGGCTGCTGGTTTTCCACT |
| **GAPDH** | AGCCACATCGCTCAGACAC | GCCCAATACGACCAAATC |
| **AARS1** | CCAGTGGCAGAAGGATGAAT | GCTTCAAGGCTTCATTCAGG |
| **CARS1** | TGAAGACCACGAAGGACTGC | GTGGGCAGACCATTTTCATCA |
| **GARS1** | TGCTGCTGAATGAGAAAGGGGAA | TACATGATCCTACCCAGGCCGA |
| **WARS1** | GTTTCCACGGACGCCCA | GACGCATTTCCCGCTTTGAG |
| **YARS1** | CAACCACGGGCAAACCACAT | CAGGTATGCGTGGAGGTCCG |
| **EIF2S2** | GATACCCAAACAGAGGAAACCC | CATCAGCTTCCAAATCCTTGTC |
| **47S rRNA** | GCTGACACGCTGTCCTCTG | ACGCGCGAGAGAACAGGAG |
| **28S rRNA** | AGAGGTAAACGGGGGGGTC | GGGGTOGGGGAGGAACGG |
| **18S rRNA** | GGGGCCCGAAGCGTTTACTTTG | CAAGAATTTCACCTCTAGCGGCGC |
| **5.8S rRNA** | GCGACGCTCAGACAGG | ACTCGGGCTCGTGCGTC |
| **5S rRNA** | GGCCATACCACCCTGAACGC | CAGCACCOGGTATTOCCAGG |

### **Supplementary Table 4**. Sequences of sgRNAs used in this study

| Gene | Forward Sequence (5' to 3') | Reverse Sequence (3' to 5') |
|---|---|---|
| **Non-targeting (NT1)** | GACGGAGGCTAAGCGTCGCAA | TTGCGACGCTTAGCCTCCGTC |
| **BRD9-g1** | CACCGCTTGACGGACAGTACCGCAG | AAACCTGCGGTACTGTCCGTCAAGC |
| **BRD9-g2** | CACCGAGGATCAACCGGTTCCTCCC | AAACGGGAGGAACCGGTTGATCCTC |
